# Emergence of short-lived meningococci causing focal epidemics can be associated with gene transfer from carriage-associated *Neisseria*

**DOI:** 10.64898/2026.09.17.752363

**Authors:** Charlene MC Rodrigues, Jay Lucidarme, Rachel Exley, Jack Llewellyn Clark, Xilian Bai, Chris Bayliss, Meera Chand, Stephen A. Clark, Cynthia Nau Cornelissen, Jeremy P Derrick, Luke R Green, Natalie Groves, Yaiza Gutierrez Vazquez, Odile B Harrison, Keith A Jolley, Samantha A McKeand, Richard Myers, Nicholas Noinaj, Kasia M Parfitt, Paolo Ribeca, Ray Borrow, Martin CJ Maiden, Christoph M Tang

## Abstract

In March 2026, an unusually large outbreak of invasive meningococcal disease (IMD) in Kent, UK, was linked to attendance at one nightclub over a single weekend. The outbreak organism was a *Neisseria meningitidis* variant belonging to the longstanding hyperinvasive genotype, cc41/44. Using genome analysis of six isolates from patients, alongside >48,000 meningococcal genomes, we investigated whether the outbreak variant had acquired traits potentially contributing to the highly invasive phenotype. The six isolates were capsular group B, sequence type (ST-)485, and essentially indistinguishable, consistent with the focal nature of the outbreak. Compared with their closest available relatives, we found changes mediated by phase variation, nucleotide variation, and horizontal gene transfer (HGT) involving adhesins, iron-acquisition systems (including Transferrin and Lactoferrin binding proteins, and FetA), and Type IV pili (Tfp), factors which affect bacteria-bacteria and bacteria-host interactions. These changes occurred in a ST-485 sub-lineage that expressed capsule at high levels and a PorA porin with a truncated surface-exposed epitope, both of which are predicted to reduce immune recognition. Donors for the HGT events were predominantly carriage-associated *N. meningitidis* and *Neisseria cinerea*. We show that meningococcal variants responsible for previous focal outbreaks have not been seen subsequently. We propose that focal outbreaks of IMD are caused by meningococcal variants that may have acquired traits from non- or less invasive organisms, but subsequently these variants disappear, as their highly invasive phenotype is inconsistent with sustained transmission. Ongoing disease surveillance alongside carriage studies are therefore essential to inform public health risk and manage epidemic IMD.

**Significance statement:** Invasive meningococcal disease (IMD), comprising sepsis and/or meningitis, is a serious life-threatening infection. IMD is usually rare, but outbreaks occur, ranging from very large epidemics to small clusters, with cases occurring over days, weeks, or months. Following an unusually large, week-long, focal outbreak in Kent, UK, in 2026, affecting 21 individuals, we investigated outbreak-associated meningococcal characteristics, comparing outbreak variant genomes with their closest available relatives. We found changes in the outbreak variants, affecting bacteria-bacteria and bacteria-host cell interactions, and iron acquisition. These traits likely resulted in an unusually high invasive potential, likely at the cost of capacity for sustained transmission in asymptomatic carriage.

## INTRODUCTION

Although *Neisseria meningitidis* is a human-adapted bacterium that asymptomatically colonizes the oropharynx, it occasionally causes invasive meningococcal disease (IMD), which is frequently fatal (1). IMD is typically caused by meningococci expressing one of six capsules, corresponding to serogroups A, B, C, W, X, and Y, that belong to a limited number of hyperinvasive genotypes, identified by MLST as clonal complexes (ccs). For example, serogroup A:cc1, A:cc5 and A:cc7 variants caused epidemics of IMD across sub-Saharan Africa (2) and C:cc1 and W:cc11 variants have caused disease outbreaks of varying scale globally (3). The UK and other high-income countries have mainly experienced endemic IMD, largely caused by B:cc41/44, B:cc32, and B:cc269 meningococci, with cases mostly occurring sporadically with occasional small clusters associated with households or schools (4–6). Larger focal outbreaks, lasting less than one week and associated with a specific transmission event, have been rare since the 1990s (7), when outbreaks caused by the ‘ET-15’ variant of C:cc11 meningococci occurred globally (8–10). These outbreaks were a major stimulus for the introduction of C conjugate polysaccharide vaccines, which effectively prevented these focal outbreaks (11).

Understanding asymptomatic carriage of the meningococcus is key to the apparently paradoxical epidemiology of IMD (12). In high income countries, meningococcal oropharyngeal carriage is highest among teenagers and young adults (13) with multiple genotypes found, most of them never or extremely rarely associated with IMD, and many expressing non-IMD associated capsules or unable to express capsule at all (*i.e.* capsule-null locus, *cnl* meningococci) (14). Meningococcal genetic and antigenic variability is driven by multiple mechanisms, including mutation, phase variation, horizontal gene transfer (HGT) with other meningococci and/or different *Neisseria* species: the meningococcus is naturally competent for DNA uptake (15, 16). Other, species of *Neisseria* represent a large gene pool accessible to meningococci conferring antimicrobial resistance and antigenic variation (17). Meningococcal diversity is consistent with the generation of multiple variants with diverse phenotypes by selective processes (18). This diversity results in the highly unpredictable nature of meningococcal outbreaks in terms of their duration and size (19).

IMD in the UK declined in the first quarter of the 21^st^ century, due to reducing meningococcal carriage rates in adolescents, a consequence of vaccine introduction and behavioral changes (20). As in many other countries, major reduction in IMD occurred during the COVID-19 pandemic, with a subsequent recrudescence, mainly due to serogroup B (21, 22). In March 2026, an exceptionally large focal outbreak occurred in Canterbury, Kent, UK, with 21 cases reported between 9^th^-16^th^ March 2026 (23). All those affected were hospitalized, nine required intensive care, and two died. Of these, 19 individuals attended the same nightclub between 5^th^-7^th^ March 2026. One further case had contact with nightclub attendees, while the other had no known contact with the nightclub; however, both lived on the same university campus as seven other cases. The UK Health Security Agency (UKHSA) sequenced the Kent outbreak variant and determined it was B:cc41/44, sequence type (ST)-485, assigned LIN code 6_0_0_0_1_27_4_18_0_0_0, using core genome MLST (24). ST-485 was the commonest ST causing serogroup B IMD in England at the time, having expanded since around 2010. The outbreak variant belonged to a larger LIN code group (6_0_0_0_1_27_4), which had a distinct PorA (P1.12-1,16-183), isolated from cases of IMD since 2020 in the UK, with additional cases in France, Spain, and Portugal (23).

We investigated the whole genome sequences (WGSs) of bacterial isolates obtained from six cases in the Kent outbreak, which were virtually indistinguishable, consistent with the point-source, focal, nature of the outbreak. The isolates were deduced to be covered by available protein-based ‘serogroup B’ vaccines. Compared to the WGSs of their closest available relatives, the outbreak isolates contained novel alleles of genes encoding adhesins, iron acquisition systems, and Type IV pili (Tfp), which mediate bacteria-bacteria and bacteria-host interactions (25). Horizontal gene transfer (HGT) involved acquisition of DNA from other serogroup B ccs, but mostly from *cnl* meningococci, carriage-associated meningococcal ccs (cc198) and the carriage-associated species, *Neisseria cinerea.* Taken together, our findings suggest focal IMD outbreaks can result from the rapid emergence of novel variants, following gene acquisition from the wider meningococcal/*Neisseria* gene pool. Retrospective analyses of meningococci from C:cc11 focal IMD outbreaks in the 1990s were consistent with our analysis.

## RESULTS

### Isolates from the Kent outbreak were highly similar and predicted to be covered by licensed ‘serogroup B’ vaccines

Of the 21 cases in the 2026 Kent outbreak, meningococci were recovered from the blood of six individuals, with the other 15 cases confirmed by PCR diagnosis. For the six isolates, single-contig, high-quality, full-length genomes were generated from short- and long-read sequences. All belonged to cc41/44 (lineage 3), ST-485, and shared the LIN code 6_0_0_0_1_27_4_18_0_0_0, consistent with these isolates possessing fewer than two loci differences in the core genome, referred to here as the ‘outbreak variant’ (*SI Appendix*, Figure S1). Comparison of the six hybrid genomes across all *Neisseria* genes catalogued in PubMLST (n=2,954) identified: i) a single allelic difference in a core gene (NEIS1452, encoding a hypothetical protein, *SI Appendix*, Table S1); ii) variation in two intergenic regions; and iii) variation in two hyper-recombinogenic loci, *pilE*/*pilS* (consistent with antigenic variation), and *mafA* in Maf Genomic Island 1 (consistent with recombination between *mafA* alleles) (26) (Figure 1*A* and *SI Appendix*, Table S2). Meningococci are subject to high-frequency phase variation through changes in repetitive DNA sequences including homopolymeric tracts (27). We detected differences among the six outbreak genomes in 11 phase variable genes (e.g., *porA*, *fetA*, *opc*, *opa*, and NEIS2780, encoding a Type I restriction-modification system, Figure 1*A*). The extensive conservation among these genomes is consistent with the focal nature of the outbreak.

**Figure 1.**
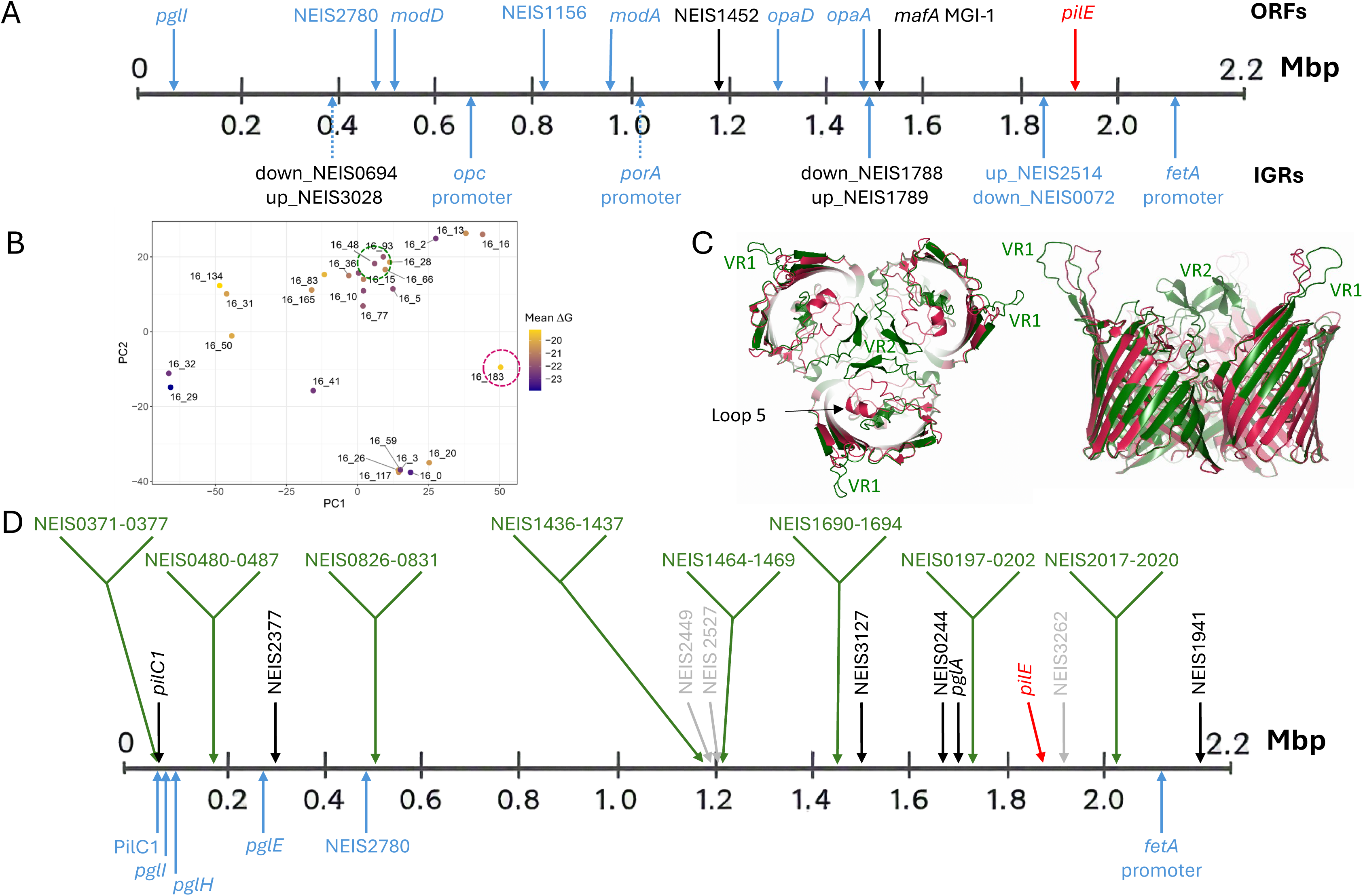
Genome comparisons of the Kent outbreak variant and its closest relatives and their PorAs. (A) Genetic differences between the six outbreak isolates by whole genome MLST. Genes with allelic variation, black; genes with phase variation, blue; genes with antigenic variation, red; variation within ORFs and in IGRs, above and below the line, respectively. (B) PCA plot comparison of predicted structures of the VR2 P16 variants of the PorA porin trimer. Points are colored by the calculated mean ΔG of the monomer interfaces in each trimer. Locations of the 16_183 and 16_48 variants are circled. (C) Orthogonal views of the PorA 16_48 (green) and 16_183 (red) structures. The locations of the VR1 and VR2 hypervariable loops are labelled for the 16_48 variant. The arrow indicates the location of loop 5 in both structures. (D) Genetic differences between the outbreak variant and closest relatives. Genes with non-synonymous changes, black; gene presence/absence, grey; genes with antigenic variation, red; genes/regions with phase variation consistently different between the 5 or 6 outbreak genomes and both closest relatives, blue. Genes involved in horizontal gene transfer events are green. NEIS nomenclature used as per PubMLST.org, gene product shown, where no gene product is shown it represents a hypothetical protein. Note, *mafA* is paralogous but only one is shown here with a difference among the outbreak isolates. Other paralogous loci not shown.

From genome data, the outbreak variant vaccine antigens were deduced to be immunologically cross-reactive with the vaccine variants in the licensed MenB vaccines (28, 29). For 4CMenB, in the outbreak variant NHBA peptide 2 was an exact match and fHbp peptide 4 was a cross-reactive match, while for MenB-fHbp, fHbp was also a cross-reactive match. The fHbp MATS relative potency (RP) for the outbreak isolates was 0.032-0.040 (mean, 0.036; median, 0.036; positive bactericidal threshold [PBT], >0.012). The corresponding values for invasive cc41/44 isolates, with fHbp peptide 4 from 2015 onwards in England range from 0.029 to 0.119 (mean, 0.051; median, 0.048). At the time of writing, only one outbreak isolate had a validated result for NHBA RP of 0.309, close to the coverage threshold PBT >0.294). The corresponding values for English invasive cc41/44 isolates, with NHBA peptide 2 from 2015 onwards, were 0.067-1.573 (mean, 0.610; median, 0.571). For the MEASURE assay for MenB-fHbp coverage, the mean fluorescence intensities ranged from 1846-2432 for the outbreak isolates, above the protective threshold for MenB-fHbp (1000), confirming the surface exposure of fHbp.

### Characteristics shared by the outbreak variant and its two closest relatives potentially impacting vaccine responses and/or invasion

Due to the high degree of similarity between outbreak isolates, one reference genome was chosen for further analysis (PubMLST id 193899, isolate id 1926231). To place the outbreak variant into its genomic context, we examined closely related genomes from ST-485, cc41/44 (lineage 3) within LIN code group 6_0_0_0_1_27_4 (45 isolates), which has caused IMD (UK, 39 cases; France, two; Austria, two; Spain and Portugal, one each). Phylogenetic analysis using cgMLST identified three close relatives, one causing an isolated case in 2022 (isolate id 22MN0044) and a two-case cluster from 2023 (isolate ids 23MN0134 and 23MN0150) from England, UK (*SI Appendix*, Figure S1); all were recovered from the blood of patients with IMD. One isolate from each cluster of IMD was chosen as the two closest available relatives (PubMLST ids 194251 and 194252).

Initially, we examined bacterial factors in the outbreak variant and its closest relatives that contribute to immune evasion. The polysialic acid serogroup B capsule confers resistance against complement-mediated killing, and thus serum bactericidal activity, an established correlate of protection against IMD (30). The capsule biosynthesis (*cssA*/*cssB*/*cssC*/*csb*) and export (*ctrA-D*) operons are expressed from divergent promoters located between the two operons (31), and polymorphisms within the 5′ untranslated region of the biosynthesis operon affects the thermal regulation and level of capsule expression. Analysis of this region suggested that the outbreak variant and its closest relatives were high-level capsule expressers (*i.e.,* igr_up_NEIS0055, allele 7), through loss of an 8bp repeat sequence compared with igr_up_NEIS0055 allele 4 (*SI Appendix*, Figure S2). Of note, 41.5% of ST-485 isolates (173 of 416 isolates) were predicted to be high capsule expressers compared with 22.6% of all other cc41/44 isolates (1061 of 4700 isolates, X2=124.6297, *df*=1, p<0.0001), which could promote the ability of ST-485 isolates to evade immune killing in the circulation.

PorA is an immunodominant integral outer membrane protein and vaccine antigen with highly antigenic, variable surface-exposed loops, termed variable regions (VRs). PorA VR2 mediates variant-specific protection conferred by OMV-based meningococcal vaccines (32). The VR2 of PorA P1.16-183 consists of only eight residues, NNNLTLVP, while other P1.16 VR2 loops contain 11 to 15 residues. To investigate the consequences of this modification, AlphaFold3 was used to predict the structures of all PorA VR2 P1.16 variants (33), which were superimposed and analyzed by Principal Component Analysis (PCA). Approximately half of the PorA VR2 variants formed a closely related cluster, while others were more dispersed; PorA VR2 P1.16-183 was well separated (circled red, Figure 1*B*). Calculation of the mean ΔG values of the monomer interfaces within trimeric PorA showed that variant P1.16-183 was predicted to be destabilized relative to most other PorA P1.16 variants. Comparison of the predicted PorA VR2 P1.16-183 structure with a member of the commonest cluster (variant 16-48) revealed that PorA VR2 P1.16-48 formed a β-hairpin, which associates with other VR2 loops on the trimer 3-fold symmetry axis (Figure 1*C*). In contrast, the shorter VR2 of PorA P1.16-183 formed contacts within the PorA barrel, displacing loop 5, indicating it is likely that this VR2 is relatively inaccessible to bactericidal antibodies.

### Differences between the outbreak variant and its closest relatives

Genomic comparisons of the outbreak variant genome to those of either or both of the two closest available relatives (PubMLST ids 194251 and 194252) identified differences in 92 genes (Table 1) and 46 IGRs (*SI Appendix*, Table S3). The 92 genetic differences included: 40 genes involved in HGT events; six genes with non-synonymous (NS) single nucleotide variants; seven genes with synonymous changes; three differences in gene presence/absence (genes of unknown function: NEIS2449, NEIS2527, NEIS3262); one gene with a single base deletion (NEIS2930); one gene with indels/multiple nucleotide variants (NEIS1426); and likely antigenic variation affecting PilE, the major pilin of Tfp (Figure 1*D*). The remaining differences were due to paralogous loci and phase variation.

**Table 1.**
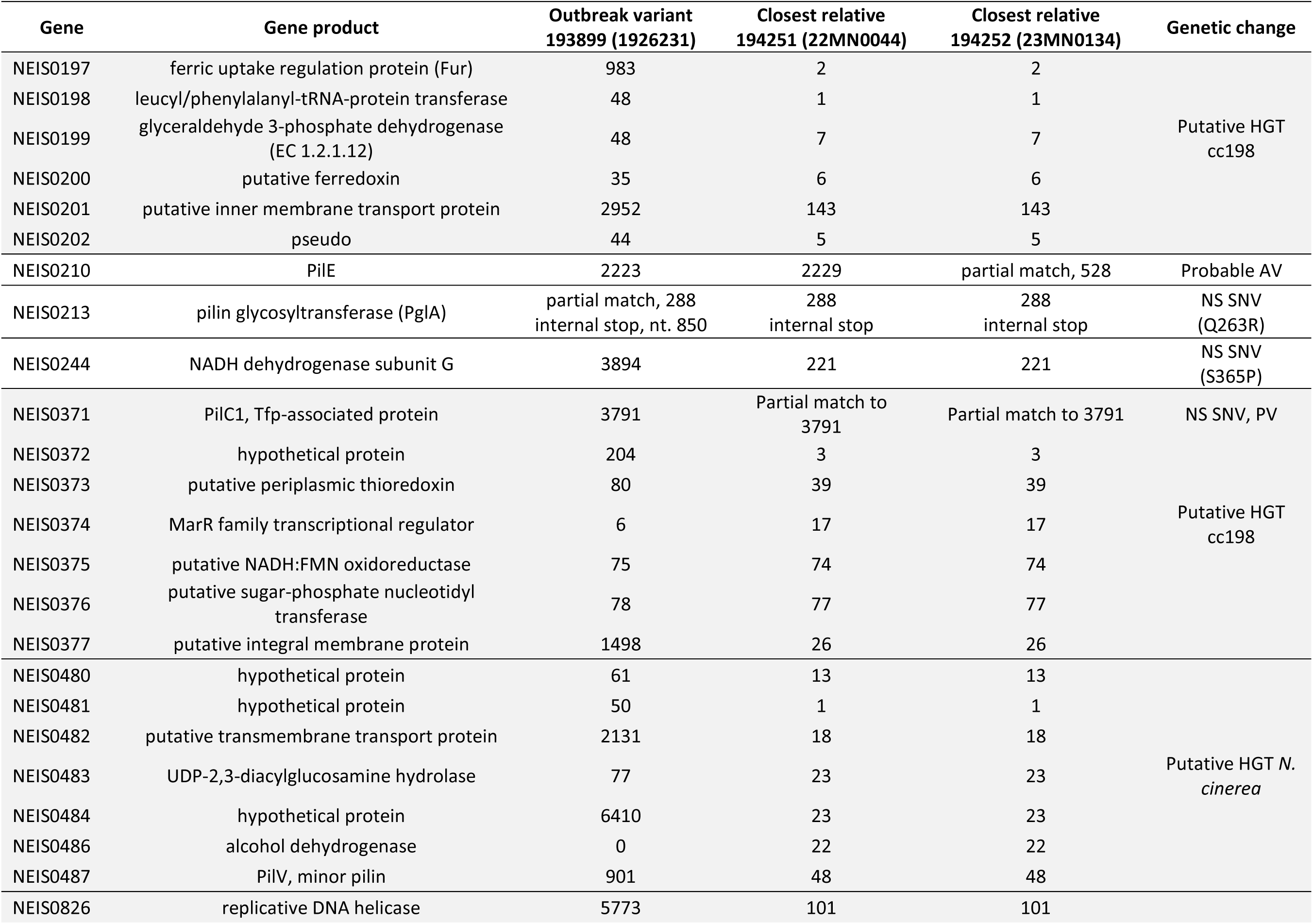
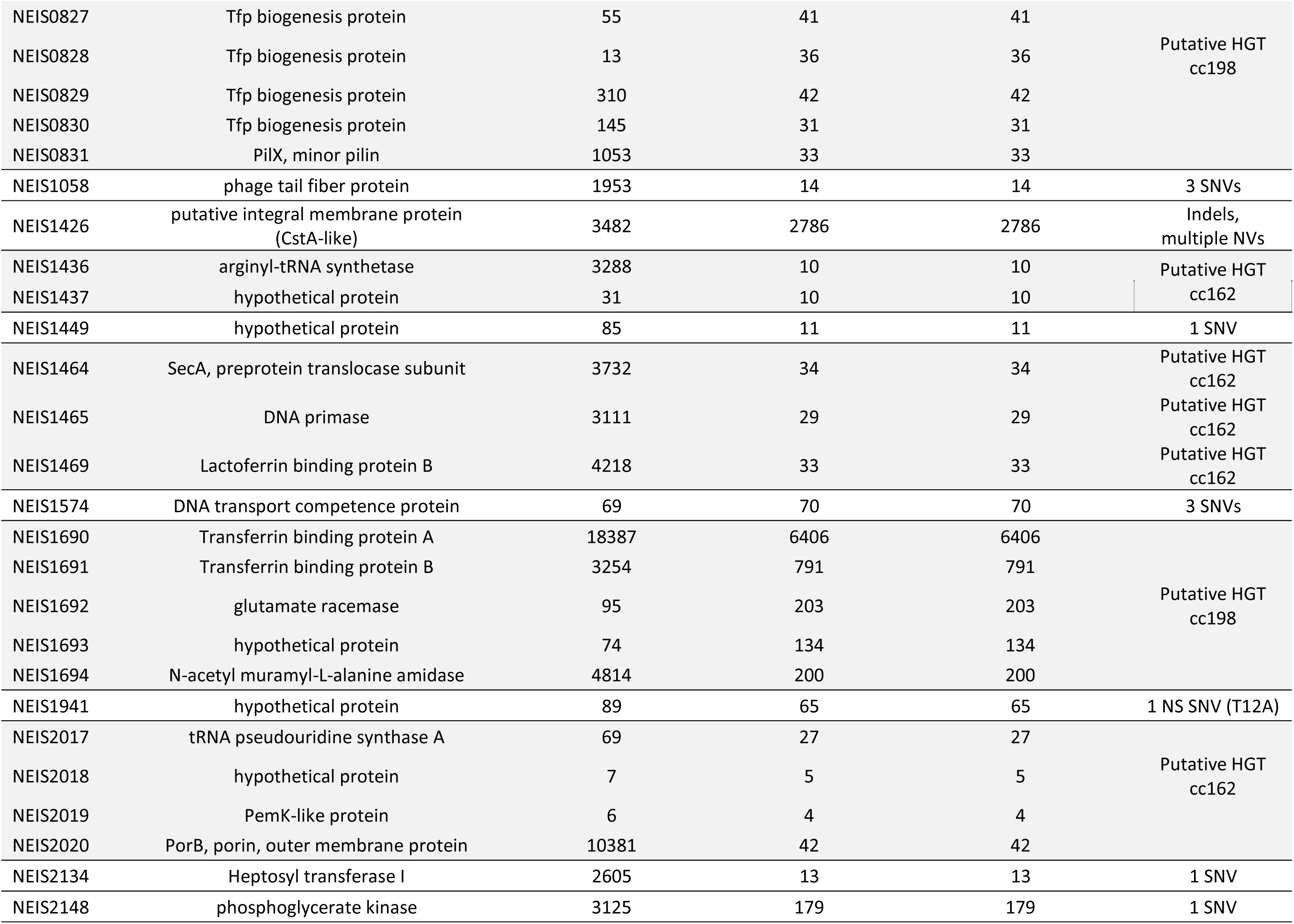
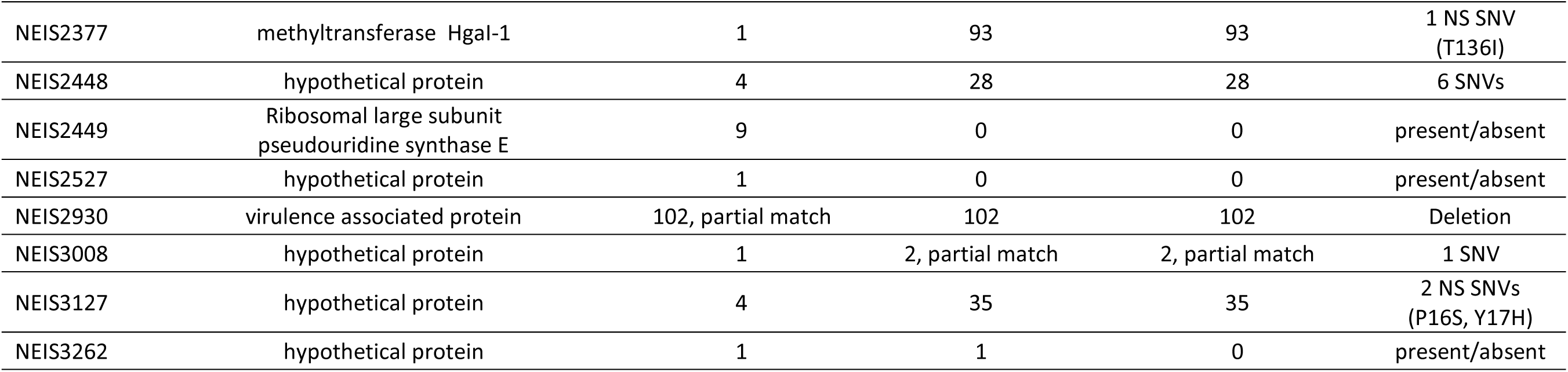
Genetic differences between the outbreak reference strain (PubMLST id:193899) and two closest relatives (PubMLST ids: 194251 and 194252) comparing all *Neisseria* genes. There were 92 changes in total, not listed are: paralogous genes and genes that differed by only phase variation in a repeat tract, for full analysis see Table S4. Amino acid changes, outbreak strains, then closest relative. Horizontal gene transfer (HGT) events shaded in grey. PV, phase variation; SNV, single nucleotide variant; NV, nucleotide variants; NS, non-synonymous. SNVs are synonymous unless otherwise mentioned.

There was a non-synonymous nucleotide change in *pglA* between the outbreak variants and its two closest relatives, although this phase-variable gene was predicted to not be translated due to the length of a polyG tract in the open reading frame (Table S4). There were non-synonymous nucleotide variations in genes encoding a subunit of NADH dehydrogenase (NEIS0244) and predicted HgaI cytosine methyl transferase (NEIS2377), and in genes of unknown function (NEIS1941, NEIS3127). Phase variation was systematically assessed in all the outbreak genomes and two closest relatives (*SI Appendix*, Figure S3 and Table S4). Six phase variable genes displayed differences between five or six of the outbreak isolates and both closest relatives (Tables S5). Two pilin glycosylation genes, *pglE* and *pglH,* were predicted OFF in the outbreak isolates and ON in both closest relatives while *pglI* had the opposite profile. As *pglA* was OFF in all isolates, the outbreak isolates likely had non-glycosylated pili, whereas the closest relatives had pili decorated with glucose (34). A Tfp-associated adhesin, PilC1, was ON in the outbreak isolates and OFF in both closest relatives while PilC2 was OFF in all isolates. FetA is phase variable due to a polyC tract in the promoter region. Depending on the number of nucleotides in the repeat, the promoter activity is weak, intermediate or strong. FetA enables the *Neisseria* species to internalize and use catechol-type siderophores produced by other bacteria (35). Of note, *fetA* was predicted to be more highly expressed in five outbreak isolates compared with its closest relatives.

Comparison with the closest available relatives identified eight putative HGT events associated with the outbreak variant (Figure 1*D*). One predicted event, NEIS2017-NEIS2020, 3.8kb, included *porB* (NEIS2020), encoding an immunodominant surface-exposed protein. The outbreak variant *porB* allele 10381, ‘porB 3-1233, was unique amongst 48,130 genomes in PubMLST. The most closely related *porB* allele 149, had one non-synonymous nucleotide substitution at nucleotide 941: this allele occurred in 293 isolates in PubMLST, with 88% in B:cc162 *N. meningitidis*, a hyperinvasive lineage (36). The combination of NEIS2017-2019 alleles predominantly occurred in cc162 isolates (253 of 241 genomes, 95.2%), consistent with this being the likely donor for this HGT event. cc162 *N. meningitidis* was a likely donor for another HGT event involving two genes, one of unknown function and the other encoding arginyl tRNA synthase (NEIS1437 and NEIS1436, respectively), with the NEIS1437 allele in the outbreak variant (33) found in 531 cc162 genomes of a total of 559 in PubMLST (95%, [accessed 17^th^ August 2026]).

### HGT events in the outbreak variant affecting iron metabolism

Three HGT events affected genes involved in iron metabolism. An ∼9.9kb region of difference between the outbreak variant and closest relatives included *tbpA* and *tbpB*. Alleles of *tbpA* and *tbpB*, involved in scavenging iron from human transferrin (hTf) were unique in the outbreak variant. The region also encoded glutamate racemase, a hypothetical protein, an amidase in peptidoglycan synthesis (NEIS1692-1694, respectively), and an additional gene of unknown function. Identification of isolates harboring identical alleles in this region (*i.e.*, NEIS1692=95, NEIS1693=74 gave 859 genomes with a defined cc/ST in PubMLST) indicated that *cnl* cc198 (accounting for 838/859, 97.3%) was a likely donor.

The outbreak *tbpA* allele (18387) had synonymous and non-synonymous (R^2^Q, outbreak to closest relatives) nucleotide differences from the closest cc198-associated allele (1354) while the outbreak variant *tbpB* allele 3254 is identical to allele 856 from a *cnl* cc198 isolate except for a single non-synonymous substitution (K^215^R, outbreak to closest relatives). Comparison of the predicted protein structure of the outbreak TbpA (allele 18387) with published structures (37, 38) and the predicted structure of the closest relative allele (allele 6406) (96% identical/97% similar) revealed high conservation of the plug and barrel domains, but multiple substitutions within the extracellular loops that interact with the C2 domain of hTf and within the loop 3 (L3) helical finger, which places a lysine side chain (K359) within the TbpA:hTf interface (Figure 2*A* and *B*). Compared to the closest relative, 75% of the sequence changes in the outbreak TbpA mapped within 15Å of the interacting interface with hTf. A three amino acid deletion in the outbreak TbpA was predicted to truncate the L3 helical finger and positions another lysine residue (K358) into the position analogous to K359 (which is substituted by N in outbreak variant) (Figure 2*C*, *D*, and *E*). The sequence changes in the L3 helix also increased the local electronegative charge along the interface with the C2 domain of hTf (Figure 2*F*, and *G*). TbpA alleles with a truncated L3 helix only occurred in 453 of 84,755 (0.5%) isolates of *Neisseria* spp. within PubMLST [accessed 8^th^ July 2026], with the majority found in *N. meningitidis* (288/453). The remainder found in non-pathogenic *Neisseria* (*e.g., Neisseria benedictiae*, *Neisseria bergeri*, *Neisseria blantyrii* and *Neisseria polysaccharea*), suggesting they act as a reservoir of Tbp diversity for the meningococcus. The 288 *N. meningitidis* isolates harbouring TbpA with a truncated L3 helix (out of 44,978 meningococci, 0.6%) were mostly in *cnl* cc198 carriage isolates (113/288). Encapsulated meningococci (menB, C and Y) carrying a TbpA truncated helix variant (116/288) were significantly more associated with IMD than those encoding a non-truncated helix (106/116, 91.4% *vs.* 17,277/22,170, 77.9%; Fisher’s exact test, *P* < 0.001, odds ratio 3.00, 95% confidence interval 1.57–5.75), suggesting that the truncated TbpAs promote IMD.

**Figure 2.**
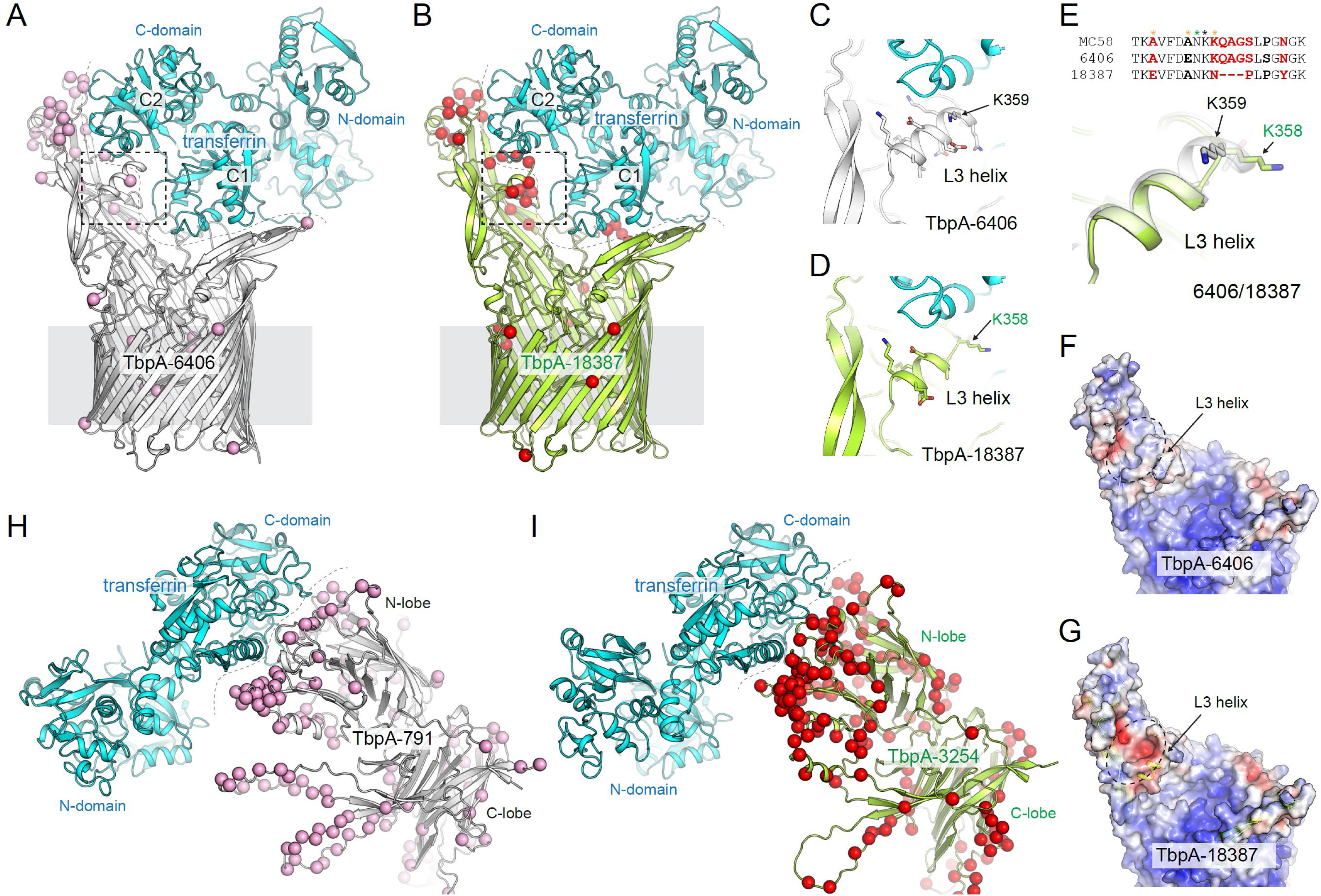
Mapping sequence changes to structural models of TbpA and TbpB. (A) Model of TbpA-6406 (grey) in complex with transferrin (cyan) with the location of residues that are different from TbpA-MC58 shown as pink spheres. 51% of the residues map to the interacting interface (grey dashed line). (B) Model of TbpA-18387 (green) in complex with transferrin (cyan) with the location of residues that are different from TbpA-6406 shown as red spheres. 75% of the residues map to the interacting interface (grey dashed line). (C) Zoomed view from panel A of the loop 3 (L3) helix of TbpA-6406 showing the residue side chains in stick representation. (D) Zoomed view from panel B of the loop 3 (L3) helix of TbpA-18387 showing the residue side chains in stick representation. (E) A sequence alignment of the L3 helix region of TbpA from MC58, 6406, and 18387. The black asterisk indicates the location of K359 (6406), the green for K358 (18387), and the orange asterisks indicate the location of residues changes affecting local electrostatic charges. Superposition of the L3 helix of TbpA from MC58, 6406, and 18387 is shown depicting the effect of the QAG deletion in 18387 in shortening the L3 helix length and the relative repositioning of K358. (F) An electrostatic surface representation (of TbpA-6406 and TbpA-18387 (G) showing the formation of a new electronegative patch (red) in proximity of the L3 helix due to the sequence changes at resides E352, A356, and N359. H) Model of TbpB-791 (grey) in complex with transferrin (cyan) with the location of residues that are different from TbpB-MC58 shown as pink spheres; 29% of the residues map to the interacting interface (grey dashed line). I) A model of TbpB-3254 (green) in complex with transferrin (cyan) with the location of residues that are different from TbpB-791 shown as red spheres; 36% of the residues map to the interacting interface (grey dashed line).

The TbpA alleles with a truncated L3 helix were consistently associated with a limited subset of TbpBs, suggesting that these transferrin receptor components are maintained together. The outbreak TbpB (allele 3254), a lipoprotein that coordinates with TbpA, has more than 30% of its total sequence change across both its N- and C-lobes compared with its closest relative (allele 791). Comparison of the predicted structure of the outbreak TbpB with published structures (37–39) and the predicted structure of the relative allele 791 (69% identical/78% similar) reveals substantial variation within the loops that interact with the C-domain of transferrin (Figure 2*H* and *I*). Compared to the closest relative, 36% of the sequence changes in the outbreak TbpB mapped within 15 Å of the interacting interface with transferrin.

A 6.4kb region spanning NEIS0197-NEIS0202 was distinct in the outbreak variant and included *fur*, encoding an iron-responsive, transcriptional regulator that controls expression of around 83 genes in *N. meningitidis* (40). Additionally, this region encodes Aat (NEIS0198), which transfers amino acids from charged tRNAs to acceptor proteins, marking them for degradation (41), glyceraldehyde 3-phosphate dehydrogenase (*gapA*, NEIS0199), a putative ferrodoxin (NEIS0200), and two genes of unknown function (NEIS0201-0202). Meningococci with the combination of alleles in this region found in the outbreak variant are predominantly *cnl* cc198 (795/830 isolates, 95.8%, therefore the likely donor); *fur* in the outbreak variant (allele 983) is a hybrid of alleles 2 (in closest relatives) and 72 (in potential donor), with the potential site of recombination between nucleotides 219-257 of the ORF. Fur was predicted to be functional in the outbreak variant as it retains domains mediating dimerisation and DNA binding, and co-ordination of Fe^2+^ (42).

While the lactoferrin binding protein A (LbpA) allele (33) was identical in the outbreak variant and closest relatives, an unpublished structure of LbpA (MC58) in complex with lactoferrin and a predicted structure of LbpA-33 reveals a shortened L3 helix finger strongly resembling that observed for TbpA-18387 (Figure *3A, B*, and *C*).

**Figure 3.**
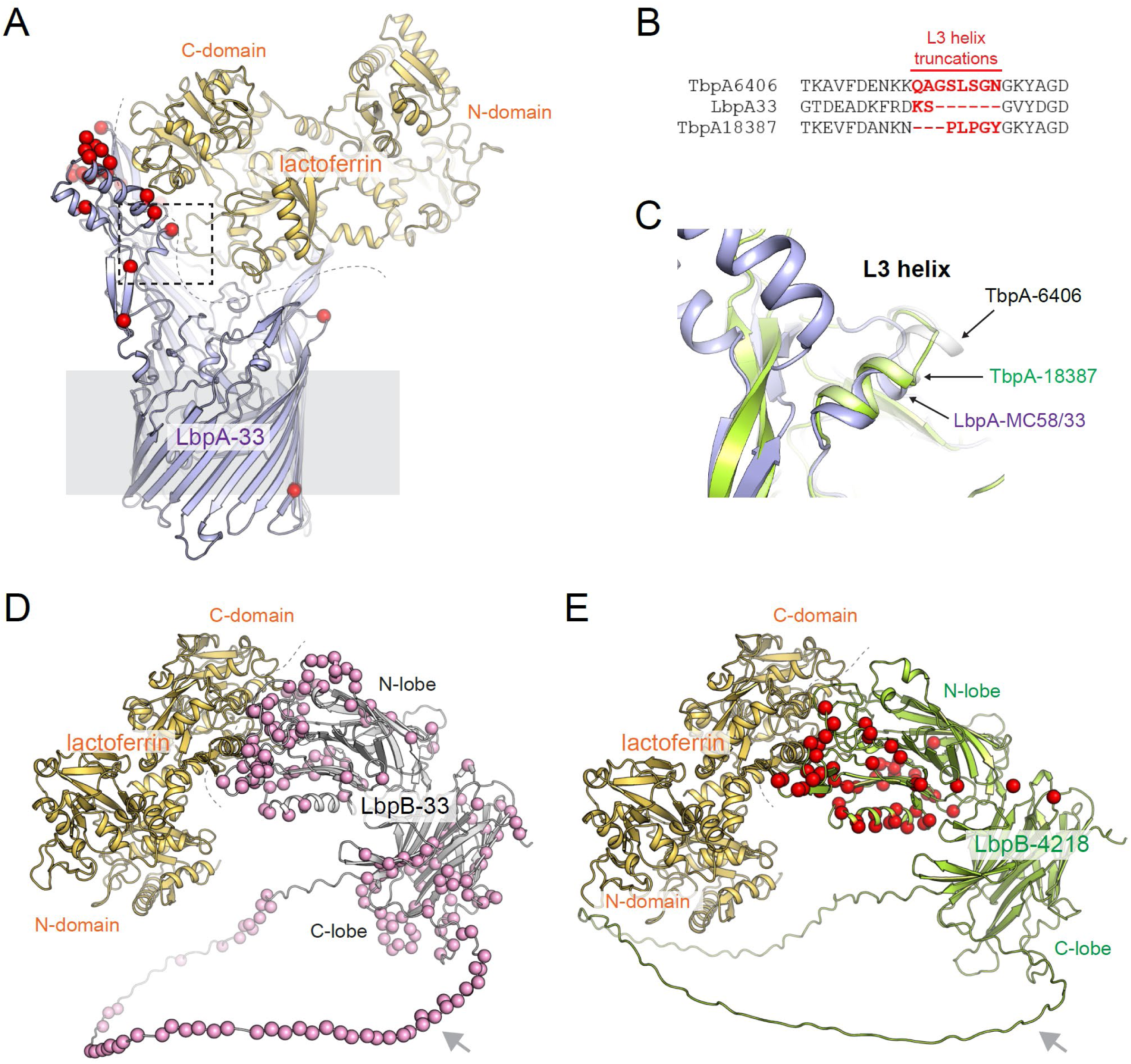
Comparison of sequence changes and structural features in models of LbpA and LbpB. (A) Model of LbpA-33 (violet) in complex with lactoferrin (gold) with the location of residues that are different from LbpA-MC58 shown as red spheres. No differences were identified between the Kent outbreak strain and its closest neighbor. (B) Sequence alignment of the L3 helix region of LbpA-33 with TbpA-6406 and TbpA-18387. LbpA-33 has a six-residue deletion within the L3 helix compared to TbpA-6406, while TbpA-18387 has a three-residue deletion. (C) A superposition of the L3 helix of TbpA from 6406 and 18387 with LbpA-MC58/33 depicting analogous shortening of the L3 helix length due to the observed sequence changes. (D) Model of LbpB-33 (grey) in complex with lactoferrin (gold) with the location of residues that are different from LbpB-MC58 shown as pink spheres; 29% of the residues map to the interacting interface (grey dashed line). The grey arrow indicates the large loop that is disordered in experimentally determined structures. E) Model of LbpB-4218 (green) in complex with lactoferrin (gold) with the location of residues that are different from LbpB-33 shown as red spheres; 57% of the residues map to the interacting interface (grey dashed line). The grey arrow indicates the large loop that is disordered in experimentally determined structures.

A further potential HGT event involved lactoferrin binding protein B (*lbpB* NEIS1469); the outbreak variant had a unique allele 4218. The first 633 nucleotides of the ORF comprising the N-lobe domain is where nucleotide variation and indels occurred compared to allele 33 (in the closest relatives) and is identical to allele 444 (Figure 3*B*), found in 15 B:cc162 isolates. The latter part of allele 4218 (C-lobe domain) is identical to the allele 33 (Figure 3*D* and *E*). Comparison of the predicted structure of the outbreak LbpB (allele 4218) with published structures (43, 44) and the predicted structure of the relative allele 33 (92% identical/94% similar) showed 57% of the sequence changes in the outbreak LbpB map within 15 Å of the interacting interface with lactoferrin. Two upstream genes that also differed between the outbreak variants and their closest relatives encode the preprotein translocase subunit and DNA primase (NEIS1464-1465, respectively). NEIS1464 allele 3732 was unique to the outbreak variant. There were eight nucleotide changes between allele 3732 and closest relative’s allele 34, located in the latter part of the sequence from nucleotide 1570 onwards: this part was identical to allele 49 found in 454 genomes, with most (343/454, 72%) in cc162 isolates, the likely donor of this HGT event. Protein structure modelling of alleles 34 and 3732 revealed identical structures with only two conservative residue substitutions (*SI Appendix*, Figure S4), so is unlikely to have functional consequences.

### HGT events affecting Tfp

Three HGT events affecting genes in Tfp biogenesis were identified (Figure 4). NEIS0371-NEIS0377 in the outbreak variant had unique alleles of genes at its boundaries (Figure 4*A*). Searches of PubMLST for genomes with identical alleles of intervening genes (NEIS0372-0376) as in the outbreak variant mostly gave *cnl* cc198 strains (333/359, 92.7%) as potential donors. PilC1 (NEIS0371) is involved with the dynamic retraction of Tfp (45) and is a potential adhesin (46), the region also includes genes encoding metabolic functions (thioredoxin, NEIS0373; NADH:FMN oxidoreductase, NEIS0375; sugar-phosphate nucleotidyl transferase, NEIS0376), a potential transcriptional regulator (NEIS0374), and a membrane protein (NEIS0376).

**Figure 4.**
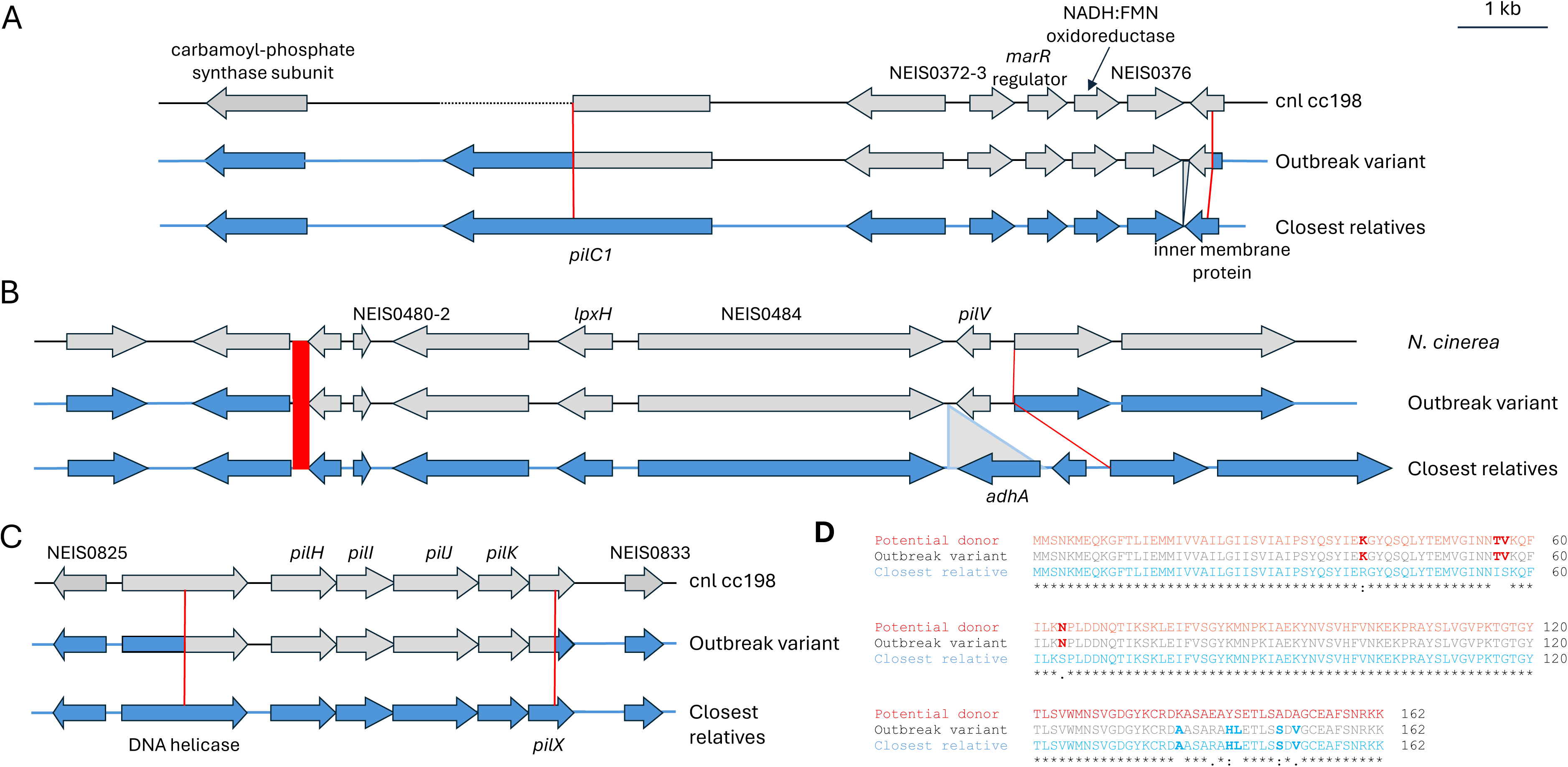
Horizontal gene transfer events involving genes involved in Tfp biogenesis and function. (A, B, C) Schematic of the alignment of genomic regions from the outbreak variant, closest relatives, and potential donors for HGT events. Open reading frames in the closest relatives and potential donors are shown as arrows filled blue and grey, respectively. NEIS numbers are shown for hypothetical proteins, and solid red lines represent predicted sites of recombination. Length of 1kb shown. (D) Clustal alignment of PilX from the outbreak variant, closest relatives, and a *cnl* cc198 meningococcal potential donor.

Another HGT event encompassed NEIS0480-NEIS0487, with the latter encoding a Tfp minor pilin, PilV, responsible for host cell signaling during attachment to epithelial cells (47). Searches using *pilV* (NEIS0487) and upstream sequences in the region yielded two *N. cinerea* isolates, consistent with inter-species recombination (Figure 4*B*). Sequences upstream of *pilV* encode hypothetical (NEIS0480, NEIS0481, NEIS0484) membrane-associated (NEIS0482) and predicted metabolic (NEIS0483) proteins. A gene for a putative alcohol dehydrogenase (NEIS0486) was absent in the outbreak variant and *N. cinerea* compared with the closest relatives. Therefore, this region from *N. cinerea* not only altered a Tfp-associated pilin but also potentially physiology and host adaptation. PilV expressed by the outbreak variant was predominately found in *N. meningitidis* (40/79, 50.6%, most commonly cc41/44 n=11, cc32 n=6, and cc11 n=5) and *N. cinerea* (27/79, 34.2%), with occasional *N. gonorrhoeae* or other *Neisseria* species (6/79, 7.6% each).

A further predicted recombination involved a Tfp-associated locus spanning NEIS0826–NEIS0831 (Figure 4*C*) with a rare *pilX* (NEIS0831) allele in the outbreak variant. PilX is a minor pilin of Tfp which is involved in bacterial aggregation and adhesion to human cells (48). Searches for isolates with identical alleles of genes within this region (*i.e*., NEIS0827-0830, inclusive) identified *cnl* cc198 isolates (268/271, 98.9%) as the likely donor. *pilH* (NEIS0827), *pilI* (NEIS0828), *pilJ* (NEIS0829) and *pilK* (NEIS0830) encode pilin-like proteins which are not components of Tfp but antagonise PilT-mediated Tfp retraction affecting Tfp dynamic behaviour (49).

Alignment of *pilX* alleles indicated that the outbreak variant allele (1053) was a hybrid generated by recombination between *pilX* in the closest relatives (allele 33) and a *cnl* cc198-associated allele (146), with the crossover between nucleotides 191 and 412 (amino acid sequence alignment, Figure 4*D*). This generated a hybrid *pilX* (allele 1053) found only in one other strain in PubMLST, a 2015 UK serogroup W:cc11 Hajj-associated pandemic isolate (PubMLST id 39411). Other isolates (n=29) expressing an identical PilX protein include serogroup A:cc1 isolates (n=23, 79%), responsible for epidemic disease in sub-Saharan Africa, and two other cc11 isolates. Isolates with a closely related PilX (*e.g.,* alleles 664 and 1860 with 97.5 and 96.9% amino acid identity, respectively) are seen in cc10217 isolates (518 and 17 isolates for alleles 664 and 1860, respectively), which caused a major outbreak in West Africa after acquiring a serogroup C capsule locus (50). Therefore, recombination generated an almost unique hybrid *pilX* in the outbreak variant, with isolates expressing closely related PilXs associated with epidemic IMD.

### Pathway enrichment analysis

Pathway analysis was used to map the transition to the outbreak variant (isolate id 1926231) from one of its closest available relatives (isolate id 23MN0134). This identified alterations in Tfp: six of the twenty genes assigned to COG category N (cell motility) differed (q = 2.0e-5), comprising the minor pilins and assembly proteins (NEIS0487, NEIS0827-NEIS0831. Two independent COG assignments, one by profile search and one by orthology, agreed closely, the latter also enriching category U, intracellular trafficking and secretion (q = 2.7e-4). The same genes were recovered through Gene Ontology as “type II secretion system complex” (q = 2.8e-4, corresponding to Tfp biogenesis, the two machineries being homologous), and an NAD(P)H oxidoreductase pair (NEIS0244, NEIS0375) was likewise enriched.

### Analysis of other focal outbreaks of IMD

Several focal IMD outbreaks occurred globally in the 1990s, caused by C:cc11 ‘ET-15’ meningococci (51). Retrospective analysis of genomes within PubMLST identified that LinCode prefix 34_0_0_0_0_0 defined cc11 ‘ET-15’ meningococci at the WGS level; these were originally characterised by Multilocus Enzyme Electrophoresis analysis (52) and sequencing of the *fumC* gene (53). This LIN Code prefix identified 2,016 ‘ET-15’ meningococci in PubMLST [accessed 17^th^ August 2026], 1,927 (95.6%) of which contained the NEIS1391(*fumC*)-50 full-length allele associated with ‘ET-15’. The oldest isolate was from the UK in 1970, 19 years before ‘ET-15’ was first described in the literature following a focal outbreak of IMD in school children in Ontario, Canada (54). Several focal outbreaks, including a large nightclub-associated outbreak in Sydney Australia, occurred over the succeeding years (55). Two UK university and one school outbreak (56) were investigated using available isolate genomes. In each case, the outbreak variants were unique (Cardiff outbreak, 1996: Unique LinCode prefix 34_0_0_0_0_0_2_38, five isolates; Southampton outbreak, 1997: Unique LinCode prefix 34_0_0_0_0_0_22, four isolates; Rotherham1999 outbreak: Unique LinCode prefix 34_0_0_0_0_0_25, two isolates), with no evidence of further spread over the succeeding 30 years. In the best characterized focal outbreak, among first year students at the University of Southampton in 1997 (8), differences between the four outbreak isolates and a closest available relative (isolate id L93/4286, PubMLST id: 644) revealed 265 differences, several of which were associated with 10 potential HGT events (Table S6). Comparisons of the Kent and Southampton outbreak WGSs with their closest available relative (*SI Appendix*, Table S7), identified that the *pilH-pilX* (NEIS0827-0831) region had changed in both outbreak variants. Of the genomes in PubMLST that share alleles of NEIS0828 and NEIS0829 with the Southampton outbreak variant (13 and 310, respectively, n=295), 92.2% are *cnl* cc198 and 6.1% are MenB, indicating that the source of the HGT was likely a non-pathogenic carried lineage.

## DISCUSSION

The outbreak in Kent 2026 represents one of the largest documented focal outbreaks of IMD associated with a single social setting. We investigated the genomic and epidemiological context of the outbreak variant, compared with its closest available phylogenetic relatives, and considered similar outbreaks. We define a focal outbreak of IMD as a temporally and spatially associated cluster of cases occurring over a short period (typically less than one week), indicating a shared acquisition source (*e.g*. a single social event) with little or no evidence of sustained onward transmission. This epidemiologic pattern is well-recognised in IMD (8–10), and differs from more protracted outbreaks, in which a defined meningococcal variant persists through community transmission, resulting in cases over extended periods, as caused by other cc41/44 variants, for example (57, 58). By combining WGS data from over 48,000 IMD isolates available in PubMLST and meningococcal carriage datasets, we were able to place the outbreak variant within a high-resolution phylogenetic framework and identified its closest available relatives. This approach illustrates the value of integrated genomic surveillance for resolving transmission and evolutionary relationships during an outbreak investigation.

To inform public health actions, the potential drivers of the Kent outbreak were considered, including: i) social behavioural and environmental factors; ii) levels of population immunity; and iii) bacterial factors, all of which contribute to meningococcal transmission and the development of invasive disease (59, 60). There did not appear to be anything exceptional about the nightclub setting or occasion responsible for the magnitude of the outbreak (23). Nevertheless, social and environmental factors are likely to have facilitated efficient transmission in a densely populated venue. Low social mixing during the COVID-19 pandemic, as well as ongoing reduction in smoking and intimate contact (20), may have led to reduced meningococcal transmission, carriage and thence low population-immunity among at the time of the outbreak (61). Consistent with this, a seroprevalence study showed that serum bactericidal activity was lower across all age groups when comparing sera from 2024-2026 against the outbreak variant with sera from 2019 against a different cc41/44 isolate, NZ 98/254 (60); however, this might also reflect differences between the target isolates, given the level of capsule production and/or truncated PorA VR2 loop in the outbreak variant.

We examined the genomic architecture of the outbreak variant to identify genetic changes within the variant that might have altered its disease-causing potential in terms of transmission dynamics and/or invasiveness (Figure 5). The outbreak variant arose from a clonal group of isolates (LIN code prefix 6_0_0_0_1_27_4, 67 allele difference threshold), comprising 45 isolates in PubMLST [accessed 7^th^ August 2026]. While these variants have been causing IMD since 2020, neither these, nor the broader B:cc41/44:ST-485 meningococcal variants, have been associated with a large focal outbreak. During the Kent outbreak, 19 of those with IMD were among 1,821 people attending a single venue over three days, an extremely high attack rate.

**Figure 5.**
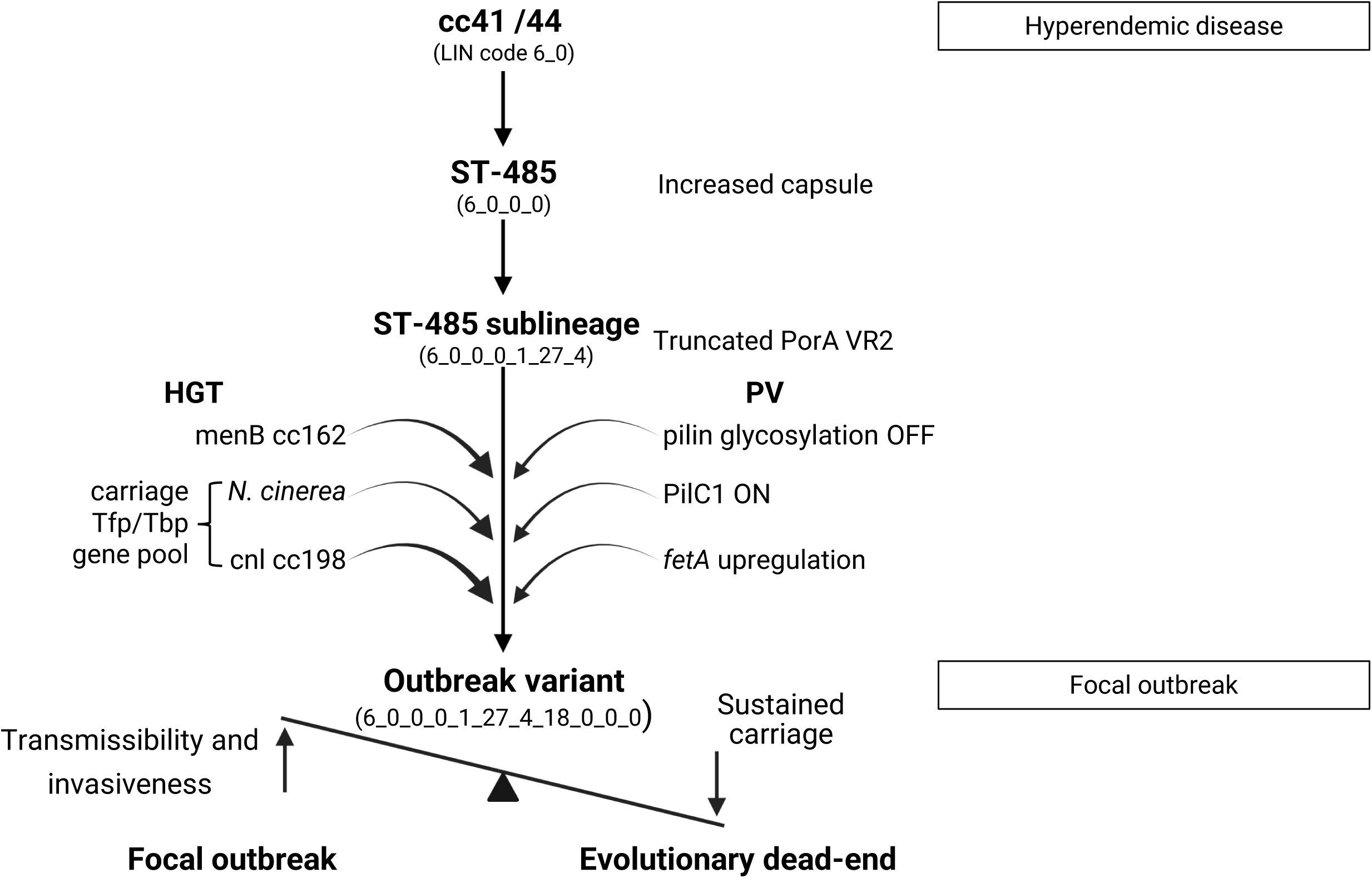
Schematic representation of the genetic changes in the Kent outbreak variant. The Kent outbreak variant is cc41/44 (LIN code 6_0), and ST-485 which has evolved increased capsule production. Within ST-485, a sublineage emerged (6_0_0_0_1_27_4) that also has a truncated PorA. Within this sublineage, the outbreak variant emerged and compared to its two closest relatives has a number of changes, including: HGT from disease-causing and carriage-associated *Neisseria;* phase variation and non-synonymous/synonymous nucleotide changes. The functional pathways that are implicated in these changes increase its invasiveness and its capacity for sustained transmission between humans. LIN - Life Identification Number, HGT – Horizontal gene transfer.

Genomic and phenotypic analyses indicate that the outbreak variant is likely susceptible to vaccine-induced immunity though NHBA and fHbp. This informed UKHSA vaccination recommendation with 4CMenB to control the outbreak in 2026 (59). PorA-mediated responses will not contribute to protection, as the outbreak variant expresses a truncated VR2 (P1.16-185 epitope), which is mismatched to the 4CMenB antigen (VR2 P1.4 epitope) (62). Immune responses and protection could also be elicited by carriage of related bacteria and mediated by meningococcal antigens, including PorA; however, with historically low carriage among UK teenagers/young adults (61) and potential VR2 inaccessibility to antibody binding, natural PorA-based host immunity against meningococci expressing PorA VR2 16-183 were likely to be negligible.

Comparison of the outbreak variant with its closest available phylogenetic relatives revealed multiple genetic differences with likely functional significance for the invasive phenotype, including HGT events, allelic variants, and phase variation (Figure 5). In the absence of contemporary carriage isolates and the three-to-four-year period since isolation of the closest relatives, it is unclear when exactly the individual genetic changes or PV states arose. Phase variation is deduced to have led to changes in expression of PilC1, an adhesin of Tfp, loss of any pilin glycosylation, and particularly of glucose moieties, and increased expression of FetA, which binds and internalises the catecolate xenosiderophores, enterobactin and salmochelin (34, 46, 63, 63).

The ability of *N. meningitidis* to acquire iron is essential for pathogenesis (64), and three regions associated with iron-acquisition differed between the outbreak variant and its closest relatives, with HGT events affecting Tbps, LbpB and Fur. The shortened L3 helix in TbpA found in the outbreak variant indicates that TbpA:hTf interactions are non-canonical; although this may alter host-pathogen interactions in ways that are not yet understood, strains expressing the shortened helix are over-represented in serogroup B disease isolates compared to those from carriage. Since this domain is both surface-exposed and critical to TbpA function (65), the loss of a few amino acids near the end of the helix in the outbreak variant could reduce its immunogenicity and/or alter iron uptake. The Lbp receptors are known to provide a competitive advantage in iron limiting environments (66), and the observed changes in the N-lobe of LbpB may enhance the capture of iron-loaded lactoferrin. Given the many substitutions in both TbpB and LbpB, both surface-exposed lipoproteins, in the outbreak variant compared to other meningococci, these changes could also contribute to immune evasion. Additionally, cc:41/44 meningococci encode both phase-variable haemoglobin-binding receptors with these receptors were switched on in all the outbreak strains. Given the central role of iron-acquisition in meningococcal virulence, such modifications could contribute to the high invasive potential of this variant.

Alterations in Tfp were another notable feature of potential HGT events detected in the outbreak variant, and affect pilin glycosylation, PilC1, *pilH-pilJ*, and the minor pilins, PilV and PilX. While the functional consequences of several of these polymorphisms remain unclear, collectively they could influence both bacterium-bacterium interactions, including biofilm formation, and bacterium-host interactions. Of particular interest, recombination at *pilX* generated a hybrid protein that is also expressed by variants associated with epidemic disease. PilX is involved in bacteria:bacteria aggregation (42). Phenotypic analysis of the only Hajj serogroup W epidemic strain with an identical *pilX* had reduced adhesion to epithelial cells and biofilm formation (67), which might impact transmission among hosts.

Parallels were evident between the focal outbreak in Kent and those seen worldwide in the 1980s and 1990s. The serogroup B Kent outbreak variant (LINCode prefix 6_0_0_0_1_27_4_18) emerged from a sublineage (‘ST-485’, LINCode prefix 6_0_0_0_1_27) of a known hyperinvasive cc (cc41/44, LinCode prefix 6_0), as was the case with the multiple variants causing serogroup C outbreaks, which were all members of the ‘ET-15’ (LINCode 34_0_0_0_0_0) part of cc11 (LINCode prefix (34_0). There is no evidence of extensive subsequent spread of the precise focal outbreak variants; although close relatives continued to cause IMD, variants associated with focal outbreaks did not. The Southampton outbreak variants (LINCode Prefix 34_0_0_0_0_0_22) were especially instructive and have been extensively investigated previously. As with the Kent outbreak, HGT with donor DNA from carriage isolates resulted in modification of Tfp-associated genes, including *pilX*, which, similar to *pilV* (involved in another HGT in the Kent outbreak), encodes a minor pilin which affects Tfp function; therefore, both focal outbreaks were caused by variants that had acquired sequences from carriage populations, either acapsulate/non-pathogenic meningococci or other *Neisseria* species. This convergence suggests that HGT with normally non-pathogenic *Neisseria* can generate phenotypes that enhance invasion. A related phenomenon was the emergence of IMD by ST-10217 meningococci, in which a large epidemic occurred in West Africa after a previously unencapsulated carriage lineage acquired a serogroup C capsule locus and a disease-associated prophage (50).

Increased invasiveness, to the levels seen in these focal outbreaks, will likely result in poor transmission. The Southampton outbreak variant and others causing focal outbreaks have not been detected in the subsequent 30 years, despite extensive global disease surveillance. As with the Kent outbreak variant, there is minimal data on carriage meningococci in the UK population since 2020 (50, 54), the time when the sublineage with a truncated PorA VR2 (LINCode prefix: 6_0_0_0_1_27_4) emerged in IMD in the UK; however, the Kent outbreak variant was not identified in these carriage studies. Given this and the observed attack rate, it is likely this variant emerged only a short period before the outbreak occurred. The outbreak variant has not been subsequently identified in the UK, potentially due to intense chemoprophylaxis (23), with only an isolated case in Austria a month after the outbreak, which was virtually identical to the outbreak variant (Figure S1). At the time of writing, there have been no further cases detected despite widespread genomic surveillance in Europe.

These data suggest a model for the evolution of the very highly invasive meningococcal phenotype where HGT between potentially invasive (encapsulated, with serogroup A, B, C, W, X, or Y capsules) and non-invasive (acapsulate) bacteria plays a role. Acquisition of traits that facilitate oropharyngeal colonization, such as enhanced inter-bacterial or host interactions, may increase local fitness, but when combined with a disease associated capsule, can promote invasion. Such combinations, if too invasive, will be evolutionarily transient, giving rise to short-lived focal outbreaks rather than sustained longer-term endemic or hyperendemic disease. This trade-off between invasiveness and sustained transmission can explain why highly invasive clusters often do not become more widely distributed or persist in meningococcal populations (Figure 5). It also explains their intrinsic unpredictability in terms of occurrence and scale (19). Understanding these relationships requires analyses of carriage and disease isolates, alongside functional studies that establish the contributions of specific genetic changes; however, the observations to date inform public health actions to continue to survey and prevent this life-threatening infection, but also highlight the need to better link genotypic and phenotypic characteristics that relate to virulence and disease risk to contextualise public health threats from genomic surveillance. With surveillance in place, robust plans for responding to outbreaks to control onward spread are essential and contact tracing, mass antibiotic prophylaxis and vaccine administration were in place within days of the first case being reported in Kent in 2026 (23). However, the biology and ecology of the meningococcus mean events like this will continue to occur, and within ST-485 at least, could develop again and as such quite unpredictably. Therefore, continued awareness of both the public and medical profession about available vaccines, their non-universal menB coverage, and the presenting signs of IMD and facilitating early recognition remain of paramount importance.

## MATERIALS AND METHODS

### Bacterial strains and genome sequencing

IMD is notifiable in England, and all isolates from culture-confirmed cases are sent to the UK Health Security Agency Meningococcal Reference Unit, this represents ∼50% of all notified cases, with the remaining PCR-confirmed (*ctrA*). Culture-confirmed IMD cases undergo routine whole genome Illumina sequencing, with genomic data published on PubMLST supporting population annotation and real-time genomic analysis (68). Routine national IMD genomic surveillance started in 2010, with 5,211 genomes from England available at the time of writing (36). Bacterial isolates (isolate ids 1926231, 1930355, 1930356, 1930357, 1934266, and 1930358) were recovered from the blood of six cases and 15 cases were confirmed by PCR. Whole genome sequences (WGS) were determined for the isolates and two closest relatives (isolate ids 22MN0044 and 23MN0134 recovered from blood) using short-(2×150 bp paired end; Illumina, NextSeq 1000 or MiSeq) and additionally long-(Oxford Nanopore Technology, ONT) read technologies. ONT libraries were prepared using the NBD114 v14 kit (SQK-NBD114-96) and loaded onto R10.4.1 flow cells and sequenced using a GridION platform. High-quality draft genome hybrid assemblies were generated (Appendix pg 1).

Genomes were annotated in PubMLST for all known meningococcal genes (69), while a custom tool was used to identify allelic variants of IGRs based on the syntenic conservation of flanking gene sequences. Core genome MLST (cgMLST) derived LIN codes were used to identify closely related meningococcal genomes among the 48,019 in PubMLST [accessed 5^th^ August 2026]. To ensure comprehensive gene coverage, PROKKA (70) and PANEROO (71) were used to identify orthologues which were compared to the NEIS locus catalogue within PubMLST, which was supplemented with the additional genes. Subsequent analyses are therefore considered to be ‘whole genome’ with respect to coding sequences. Genome comparator using cgMLST (n=1329) and whole-genome MLST (wgMLST) (n=2954 loci) was used on PubMLST.org. For comparing genomes, the first Kent outbreak isolate submitted to PubMLST (isolate id 1926231; hybrid genome PubMLST id 193899) was used as the reference outbreak strain.

### Analysis of potential phase variation and HGT events

Identification and analysis of phase-variation was performed using PhasomeIt v1.1.5 (72). Phase-variable repeat tracts were detected using the following minimum thresholds for repeat number based on tract size and type: polyC or G - 9; polyA or T - 11; di- and tri-nucleotide - 8; tetra- and penta-nucleotide - 5; hexa-nucleotide or greater – 3. A subset of tract lengths was checked for agreement with raw read data.

Potential HGT events were identified as: i) genome regions with >1 allele difference over a block of syntenic loci; or ii) alleles which differ by > 20 nuceleotides, for which a corresponding region was found in another *Neisseria* genome in PubMLST. The allelic profile of syntenic loci that differed from the closest relative was used to search *Neisseria* isolates in PubMLST using the ‘locus =’ with the ‘AND’ function. Potential recombination sites were identified with Clustal Omega (73) and SnapGene (74). Sequences corresponding to potentially horizontally acquired loci in the outbreak strain and flanking loci were extracted from PubMLST and aligned to the corresponding sequence from the closest available relative, and potential donor.

### Assessment of vaccine coverage by antigen expression

Meningococcal Antigen Typing System (MATS) was used to estimate variant coverage by 4CMenB (Bexsero, GlaxoSmithKline) (62). The Meningococcal Antigen Surface Expression (MEASURE) assay was used to estimate variant coverage by MenB-FHbp (Trumenba, Pfizer) (75). Meningococcal Deduced Vaccine Antigen Reactivity (MenDeVAR) Index and Genetic Meningococcal Antigen Typing System (gMATS) were used to deduce cross-reactivity from sequence data (28, 29).

### Protein modelling

Structures were predicted using AlphaFold3 and exported as mmCIF files (33). Structures were converted to PDB format using Gemmi (76) and analysed in R (77). For PorA, as all three chains within each model were identical, only chain A was retained for downstream analysis.

Coordinates for backbone atoms (N, Cα, C and O) were extracted and structures aligned using MUSCLE, followed by least-squares superposition of equivalent backbone positions. A structure-informed multiple sequence alignment was generated from the superposed models and used to identify homologous residue positions across variants. Principal component analysis (PCA) was performed on the Cartesian coordinates of aligned backbone atoms using the Bio3D function pca.xyz. Mean ΔG values of the monomer interfaces were calculated using PISA.

Similarly, the outbreak strain sequences were used to predict structural models of the Tbps and Lbps in complex with human transferrin and lactoferrin, respectively, and visualised, and aligned to unpublished (LbpA-MC58 in complex with lactoferrin; Yadav, Dubey, Noinaj) and published structures (37–39, 43, 44) using PyMOL (Schrödinger LLC) (78).

### Pathway enrichment analysis

Protein sequences were translated from each genome’s own GFF3 annotation, all three produced by the PubMLST pipeline (69), using the bacterial genetic code with canonical start codons read as methionine, and high-quality, hybrid ONT/Illumina genomes of a close relative of outbreak variant (PubMLST id 194252) and the outbreak variant (PubMLST id 193899).

Predicted proteomes were compared locus by locus; a locus contributed to a comparison only where both genomes carried a complete, high-quality coding sequence, excluding: (i) features flagged as incomplete; (ii) of length not divisible by three; (iii) carrying an internal stop; or (iv) containing ambiguous bases. Two classes of difference were excluded from both comparisons to keep them equivalent: presence/absence, and copy-number changes. The latter was done because all 55 such differences originated from copies lost in the ancestral genome alone, a one-directional pattern consistent with the different quality of the assemblies rather than with biology.

The background for both tests was the set of 2008 loci carrying a complete, high-quality coding sequence in all three genomes, that is, the genes that could have differed in either comparison; substituting the union of the three genomes yielded equivalent results. Functional terms were obtained by three independent routes, kept separate because their coverage differs, and each recovers terms the others miss. Gene Ontology (79) and KEGG (80) terms were transferred from *N. meningitidis* MC58 through locus aliases (1487 of 2008 loci); COG categories were assigned de novo from the isolates’ own proteins by RPS-BLAST against the COG/CDD profile database (81, 82), as implemented in COGclassifier v2.0.0 (1574 loci); and Gene Ontology, KEGG and COG were assigned de novo by orthology using eggNOG-mapper v2 (83) against eggNOG 5.0 (84) (1762 loci), with GO terms propagated up the “is_a” and “part_of” hierarchy. Together, the three routes annotated 1821 of the 2008 background loci (91%). Over-representation was assessed within each scheme by the hypergeometric test with Benjamini-Hochberg correction (85), considering terms carrying at least two genes; restricting each background to genes annotated in the scheme under test led to the same conclusions.

## Author Contributions

Conceptualization: CMCR, JL, RE, MC, RM, RB, MCJM, CMT. Methodology: CMCR, JL, RE, JLC, CB, MC, CNC, JPD, LRG, YGV, OBH, KAJ, SAM, RM, NN, KMP, RB, MCJM, CMT. Software: JLC, CB, JPD, LRG, NG, YGV, OBH, KAJ, SAM, RM, NN, KMP, PR, MCJM, CMT. Validation: CMCR, JL, RE, JLC, XB, CB, SC, CNC, JPD, LRG, YGV, OBH, KAJ, SAM, RM, NN, PR, RB, MCJM, CMT. Formal analysis: CMCR, RE, JLC, CB, CNC, JPD, LRG, NG, YGV, OBH, KAJ, SAM, RM, NN, PR, MCJM, CMT. Investigation: JL, RE, JLC, XB, CB, SC, CNC. Resources: JL, MC, SC, RM, RB, MCJM, CMT. Data curation: CMCR, JL, RE, JLC, CB, LRG, YGV, OBH, KAJ, SAM, RM, RB, MCJM, CMT. Writing – original draft: CMCR, JL, JLC, CB, JPD, LRG, YGV, SAM, NN, PR, RB, MCJM, CMT. Writing – review & editing: CMCR, JL, RE, JLC, XB, CB, MC, SC, CNC, JPD, LRG, NG, YGV, OBH, KAJ, SAM, RM, NN, KMP, PR, RB, MCJM, CMT. Visualization: CMCR, JLC, CB, CNC, JPD, NG, SAM, NN, PR, MCJM, CMT. Supervision: CMCR, JL, RE, RB, MCJM, CMT. Project administration: CMCR, RB, MCJM, CMT. Funding acquisition: MCJM, CMT, NN, CNC

## Classification

Biological Sciences, Microbiology

## Data sharing

All hybrid genomes used in this study are publicly-available on PubMLST.org under the PubMLST ID given in the paper. The isolate numbers provided can be used to retrieve assembled genomes that were generated using Illumina read data only.

## Acknowledgements

Thank you to staff at the UKHSA Meningococcal Reference Unit for preliminary culture and typing and staff at Pathogen Genomics Sequencing Lab - Manchester for performing DNA extraction and genome sequencing. Materials and support for the MATS assay were provided by GlaxoSmithKline Biologicals SA. GlaxoSmithKline Biologicals SA had no role in data collection, analysis, interpretation, writing of the manuscript or the decision to submit for publication. Thank you to Helen Campbell and Shamez Ladhani for providing epidemiology data.

