## Supplemental data for "Emergence of short-lived meningococci causing focal epidemics can be associated with gene transfer from carriage-associated *Neisseria*"

**Appendix**

**Methods**

**Hybrid genomes generation**

Dorado (version 0.9.1) (<https://github.com/nanoporetech/dorado>) was used to remove adapters and barcode sequences from reads. Reads were trimmed with Chopper (version 0.9.0) (<https://academic.oup.com/bioinformatics/article/39/5/btad311/7160911?login=true>) to remove low quality (PHRED Q-score <10) and short reads (<150bp). Samples that contained <50 reads were removed from downstream analysis.

Reads were randomly down-sampled with Rasusa (version 2.1.0) (<https://joss.theoj.org/papers/10.21105/joss.03941>) to enable fast and effective initial assembly with Raven (version 1.8.3) (<https://www.nature.com/articles/s43588-021-00073-4>). The initial assembly was used to estimate the genome size and expected coverage of the reads, with this information being fed into Autocycler (version 0.5.1) (<https://academic.oup.com/bioinformatics/article/41/9/btaf474/8242761>). Reads were first subsampled by Autocycler into four minimally overlapping subsets of reads. Each of these subsets were then assembled by Canu (version 2.3) ([https://doi.org/10.1101/gr.215087.116](https://pmc.ncbi.nlm.nih.gov/articles/PMC5411767/)), Flye (version 2.9.6) (<https://www.nature.com/articles/s41587-019-0072-8>), Miniasm (version 0.3) (<https://academic.oup.com/bioinformatics/article/32/14/2103/1742895>) and Myloasm (version 0.2.0) (<https://www.biorxiv.org/content/10.1101/2025.09.05.674543v1>). A consensus weighting of two was given to Canu and Flye assemblies. A cluster weighting of two was given the Myloasm assemblies where the sequence was not duplicated and depth was in the single digits. Consensus assemblies were then generated following Autocycler default settings. Any Autocycler assembly fails resulted in a “fallback” assembly, where Flye alone was used to generate an assembly. All resulting assemblies were reoriented with Dnaapler (version 1.2.0) (<https://joss.theoj.org/papers/10.21105/joss.05968>) to ensure a consistent starting point and sense for each circular contig. Concurrently to Autocycler, Plassembler (version 1.8.0) (<https://academic.oup.com/bioinformatics/article/39/7/btad409/7208863>) was run on a Rasusa down-sampled set of reads to 60x coverage to detect the presence of any small plasmids that may otherwise have been missed. Any resulting plasmids that were identified by Plassembler were reoriented with Dnaapler and dereplicated with skDER (version 1.3.4) (<https://pubmed.ncbi.nlm.nih.gov/40637375/>) using a percentage identity and aligned fraction cut-off of 98 and 90 (percent) respectively. SkDER runs on top of the Skani (version 0.3.1) software (<https://www.nature.com/articles/s41592-023-02018-3>), where options for working with small genomes and plasmids were enabled (-c 30, -m 200). Kraken2 (version 2.1.5) (<https://link.springer.com/article/10.1186/s13059-019-1891-0>) was used to confirm that the sample was indeed bacterial, with this information used to select the appropriate polishing model for aligning and polishing with Dorado. ONT assemblies were then polished using Illumina reads with one iteration of Polypolish version 0.6.0 followed by an iteration of Pypolca version 0.3.1.

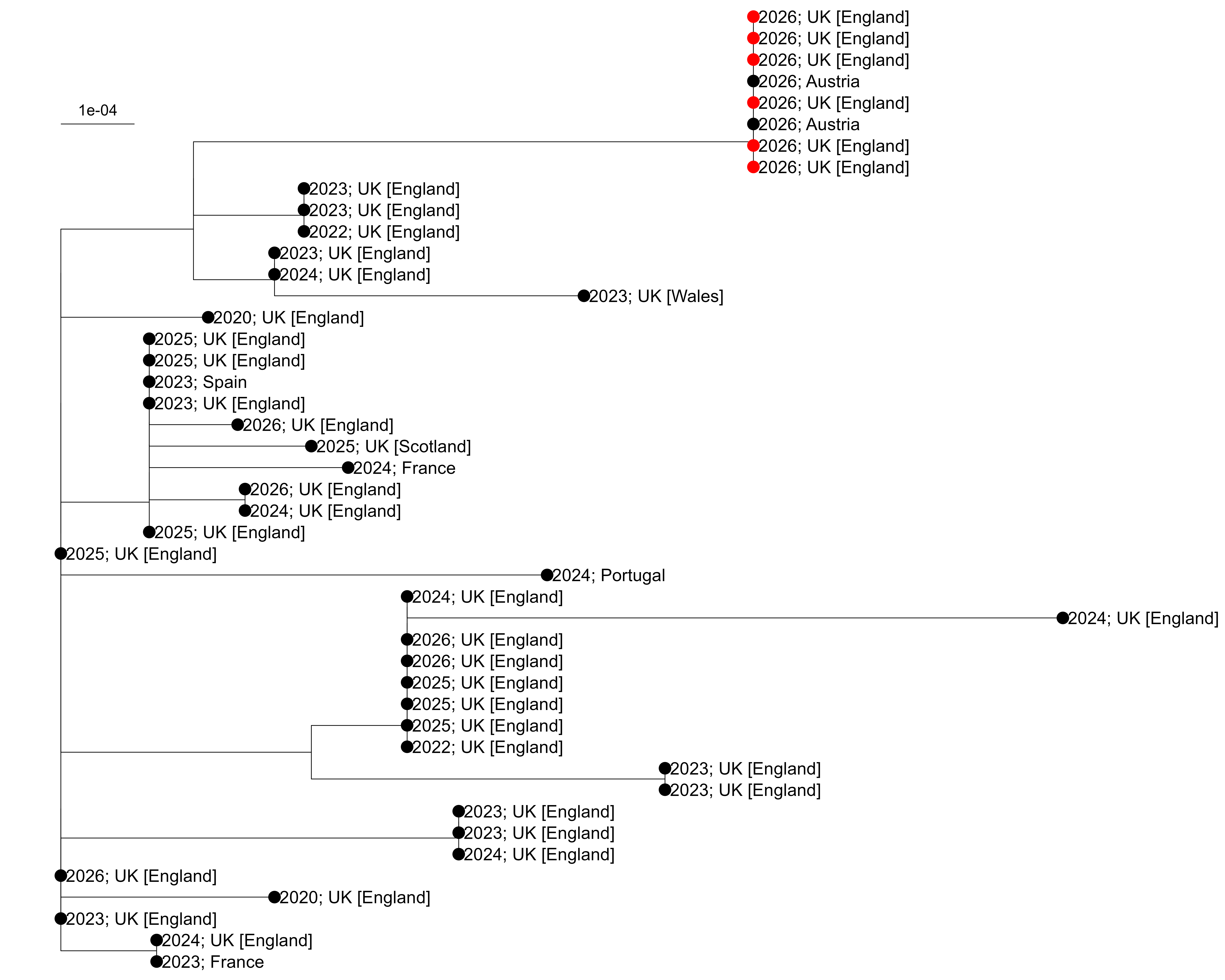

**Figure S1:** Whole genome multilocus sequence typing neighbour-joining tree generated from sequences with ClustalW containing all sequences within LIN code group 6_0_0_0_1_27_4 n = 45 isolates). The sequences from the outbreak variant from Kent are shown in red. All sequences are labelled with year and country of collection. Generated with sequence data available in PubMLST using MicroReact (https://microreact.org/project/3escbvUje3os58CidXmaQJ-bigsdb3458369907286637801460)

**Table S1:** Genetic allelic differences across the core genome (n=1329) between the six outbreak genomes (PubMLST ids:193899; 19390; 194248; 194249; 194250; 195530, isolate identifiers are shown in brackets after each one).

| **Locus** | **Product** | **193899 (1926231)** | **193900 (1930355)** | **194248 (1930356)** | **194249 (1930357)** | **194250 (1934266)** | **195530 (1930358)** |
| --- | --- | --- | --- | --- | --- | --- | --- |
| NEIS1452 | Hypothetical protein | 1964 | 1964 | 1964 | 105 | 1964 | 1964 |

**Table S2.** Gene and intergenic allelic differences across the whole genome (n=2,954) between the six outbreak isolates (PubMLST ids:193899, 19390, 194248, 194249, 194250; 195530; isolate identifiers are shown in brackets). There were 23 changes in total (16 in ORFs and seven in IGRs), not listed in the table are: (i) six genes that are probable paralogues with no clear evidence of differences between the isolates: NEIS0041, NEIS0596, NEIS1574, NEIS1789, NEIS1880, and NEIS1996; (ii) seven phase variable genes where variation occurs only in the repeat tract: NEIS0380 (*pglI*), NEIS1156, NEIS1310 (*modA*), *opaA*, *opaD*, NEIS2364 *(modD),* and NEIS2780; and (iii) four phase variable promoter regions of NEIS1364 (*porA*), NEIS1963 (*fetA*), NEIS2198 (*opc*) and an intergenic region upstream of NEIS2514. Numbers indicate the alleles in PubMLST for each isolate.

| **Locus** | **Product** | **193899 (1926231)** | **193900 (1930355)** | **194248 (1930356)** | **194249 (1930357)** | **194250 (1934266)** | **195530 (1930358)** |
| --- | --- | --- | --- | --- | --- | --- | --- |
| NEIS0210 | PilE | 2223 | 2226 | 2227 | 2223 | 2228 | 2235 |
| NEIS1452 | Hypothetical protein | 1964 | 1964 | 1964 | 105 | 1964 | 1964 |
| NEIS1789 | MafA1 lipoprotein | 29; 32 | 29; 32 | 29; 32 | 29; 32 | 29; 32 | 29; 32 |
| igr_down_NEIS1788_  up_NEIS1789 |  | 1 | 1 | 1 | 1 | 2 | 1 |
| igr_down_NEIS0694_  up_NEIS3028 |  | 1 | 2 | 1 | 1 | 1 | 1 |

**Table S3:** Intergenic differences between the outbreak variant (PubMLST id:193899) and two closest relatives (PubMLST ids: 194251 and 194252). A custom tool was used to identify allelic variants of IGRs based on the syntenic conservation of flanking gene sequences.

| **IGR** | **Kent outbreak strain**  **193899 (1926231)** | **Closest relative**  **194251 (22MN0044)** | **Closest relative**  **194252 (23MN0134)** | **Genetic change** |
| --- | --- | --- | --- | --- |
| igr_down_NEIS3006_down_NEIS0371 | 1 | 2 | 2 | 1 SNV |
| igr_up_NEIS0371_down_NEIS0367 | 1 | 2 | 2 | 6 SNVs |
| igr_up_NEIS0372_up_NEIS0373 | 1 | 2 | 2 | 3 SNVs |
| igr_down_NEIS0373_up_NEIS3008 | 1 |  |  |  |
| igr_down_NEIS0374_up_NEIS0375 | 1 | 2 | 2 | 1G insertion |
| igr_down_NEIS0376_down_NEIS0377 | 1 |  |  |  |
| igr_up_NEIS0480_up_NEIS0481 | 1 | 2 | 2 | 3 SNVs |
| igr_down_NEIS0481_down_NEIS0482 | 1 | 2 | 2 | 6 SNVs |
| igr_up_NEIS0482_down_NEIS0483 | 1 | 2 | 2 | 3 SNVs |
| igr_up_NEIS0483_up_NEIS0484 | 1 | 2 | 2 | 4 SNVs, 1A insertion |
| igr_down_NEIS0484_down_NEIS0487 | 1 |  |  |  |
| igr_up_NEIS0487_up_NEIS0488 | 1 | 2 | 2 | 8 SNVs |
| igr_down_NEIS3265_up_NEIS2786 | 1 | 2 | 2 | CAAA insertion |
| igr_down_NEIS2786_up_NEIS2563 | 1 | 2 | 2 | 1G deletion |
| igr_up_NEIS0651_up_NEIS0652 | 1 | 2 | 2 | 1 SNV |
| igr_down_NEIS0694_up_NEIS3028 | 1 | 2 | 2 | CTCATTT deletion |
| igr_down_NEIS0826_up_NEIS0827 | 1 | 2 | 2 | 1 SNV |
| igr_up_NEIS2553_down_NEIS2552 | 1 | 2 | 2 | 1 SNV |
| igr_up_NEIS1145_up_NEIS1146 | 1 | 2 | 2 | 2 SNVs |
| igr_down_NEIS1278_up_NEIS1279 | 1 | 2 | 2 | 1 SNV |
| igr_down_NEIS2448_up_NEIS2527 | 1 | 2 | 2 | 10 SNVs, 1C insertion |
| igr_down_NEIS2527_up_NEIS2449 | 1 |  |  |  |
| igr_down_NEIS2449_up_NEIS1428 | 1 |  |  |  |
| igr_down_NEIS1436_up_NEIS1437 | 1 | 2 |  |  |
| igr_down_NEIS1437_down_NEIS1438 | 1 |  |  |  |
| igr_down_NEIS1464_up_NEIS1465 | 1 | 2 | 2 | 2 SNVs |
| igr_up_NEIS1469_down_NEIS1471 | 1 |  |  |  |
| igr_up_NEIS1688_up_NEIS1689 | 1 | 2 | 2 | 12 SNVs |
| igr_down_NEIS1689_up_NEIS3127 | 1 | 2 | 2 | 7 SNVs, 1A deletion |
| igr_down_NEIS3127_down_NEIS1690 | 1 |  |  |  |
| igr_up_NEIS1690_down_NEIS1691 | 1 | 2 | 2 | 1 SNV |
| igr_up_NEIS1691_down_NEIS1692 | 1 |  |  |  |
| igr_up_NEIS1692_up_NEIS1693 | 1 | 2 | 2 | 7 SNVs, 1G insertion |
| igr_up_NEIS1859_down_NEIS3148 | 1 | 2 |  |  |
| igr_up_NEIS1879_down_NEIS1882 | 1 | 2 | 3 | 19 SNVs |
| igr_up_NEIS0201_up_NEIS0200 | 1 | 2 | 2 | 1 SNV |
| igr_down_NEIS0200_up_NEIS0199 | 1 | 2 | 2 | 5 SNVs, 1A deletion |
| igr_down_NEIS0199_down_NEIS0198 | 1 | 2 | 2 | 1 SNV, 7 nt deletion, 39 nt insertion |
| igr_up_BACT000032_down_BACT000010 | 1 | 2 | 2 | 3 SNVs |
| igr_up_NEIS0041_down_NEIS2981 | 1 | 2 | 2 | 3 SNVs |
| igr_down_NEIS0210_up_NEIS0001 | 1 |  | 2 |  |
| igr_down_NEIS2142_up_NEIS2141 | 1 | 2 | 2 | 1T deletion |
| igr_up_NEIS2988_down_NEIS2019 | 1 |  |  |  |
| igr_up_NEIS2018_down_NEIS2017 | 1 | 2 | 2 | 4 SNVs |
| igr_up_NEIS2017_down_NEIS3256 | 1 | 2 | 2 | GTT insertion |
| igr_down_NEIS1997_up_NEIS1996 | 1 | 2 | 2 | 2 SNVs |

**Table S4:** For phase variable (PV) genes across six outbreak genomes (PubMLST ids:193899; 19390; 194248; 194249; 194250; 195530) and their closest relatives (PubMLST ids: 194251 and 194252). ON and OFF states for genes with PV tracts within the coding region are in green and red text, respectively. Where they are known, expression states of loci with PV tracts in IGRs are boxed orange, yellow and dark green for low, intermediate and high expression, respectively. Other genes with PV tracts in an IGR are uncoloured.

| **Gene** | **Gene id** | **193899 (1926231)** | **193900 (1930355)** | **194248 (1930356)** | **194249 (1930357)** | **194250 (1934266)** | **195530 (1930358)** | **194251 (22MN0044)** | **194252 (23MN0134)** | **Tract variation among outbreak isolates^1^** | **Tract variation between outbreak and neighbours^2^** | **Expression different between outbreak and both neighbours^3^** |
| --- | --- | --- | --- | --- | --- | --- | --- | --- | --- | --- | --- | --- |
| Pilus assembly and glycosylation | | | | | | | | | | | | |
| *pilC1* | NEIS0371 | G10 | G10 | G10 | G10 | G10 | G10 | G12 | G12 | No | Yes | Strong (6/6) |
| *pilC2* | NEIS0033 | G11 | G11 | G11 | G11 | G11 | G11 | G11 | G11 | No | No | N/A |
| *pglA* | NEIS0213 | G9 | G9 | G9 | G9 | G9 | G9 | G9 | G9 | No | No | N/A |
| *pglH* | NEIS0400 | C11 | C11 | C11 | C11 | C11 | C11 | C10 | C10 | No | Yes | Strong (6/6) |
| *pglI* | NEIS0380 | G13 | G13 | G13 | G12 | G13 | G13 | G11 | G12 | Yes | Yes | Strong (5/6) |
| *pglG* | NEIS0401 | C11 | C11 | C11 | C11 | C11 | C11 | C11 | C11 | No | No | N/A |
| *pglE* (AAACAAC) | NEIS0568 | 9 | 9 | 9 | 9 | 9 | 9 | 8 | 11 | No | Yes | Strong (6/6) |
| Adhesins | | | | | | | | | | | | |
| *opaA^4^* |  | 8 | 6 | 8 | 8 | 9 | 7 | 9 | 6 | Yes | Yes | Weak (3/6) |
| *opaB^4^* |  | 10 | 10 | 10 | 10 | 10 | 10 | 12 | 16 | No | Yes | N/A |
| *opaD^4^* |  | 23 | 22 | 14 | 22 | 22 | 22 | 16 | 19 | Yes | Yes | N/A |
| *opaJ^4^* |  | 10 | 10 | 10 | 10 | 10 | 10 | 8 | 7 | No | Yes | N/A |
| *opc* | NEIS2198 | **C15** | **C15** | **C15** | C14 | **C15** | C14 | C11 | C11 | Yes | Yes | Weak (4/6) |
| RMS, other regulatory | | | | | | | | | | | | |
| *modB* (CCCAA) | NEIS1194 | 28 | 28 | 28 | 28 | 28 | 28 | 22 | 22 | No | Yes | N/A |
| *modA* (AGCC) | NEIS1310 | 28 | 31 | 29 | 29 | 29 | 28 | 30 | 11 | Yes | Yes | Weak (3/6) |
| *modD* (AACCG) | NEIS2364 | 9 | 9 | 8 | 8 | 9 | 9 | 11 | 11 | Yes | Yes | N/A |
| Type I RMS, specificity subunit | NEIS2780 | G9 | G9 | G9 | G10 | G9 | G9 | G10 | G8 | Yes | Yes | Strong (5/6) |
| Iron acquisition | | | | | | | | | | | | |
| *fetA* | NEIS1963 | C10 | C10 | C11 | C10 | C10 | C9 | **C8** | C9 | Yes | Yes | Strong (5/6) |
| *hpuA* | NEIS1946 | G10 | G10 | G10 | G10 | G10 | G10 | G10 | G10 | No | No | N/A |
| *hmbR* | hmbR | G9 | G9 | G9 | G9 | G9 | G9 | G9 | G9 | No | No | N/A |
| Membrane transport | | | | | | | | | | | | |
| *porA* | NEIS1364 | G11 | G11 | G12 | G12 | G12 | G12 | G11 | G11 | Yes | Yes | N/A |
| peptidase | NEIS2992 | C9 | C9 | C9 | C9 | C9 | C9 | C9 | C9 | No | No | N/A |
| Immune evasion and survival | | | | | | | | | | | | |
| *nalP* | NEIS1943 | C10 | C10 | C10 | C10 | C10 | C10 | C9 | C10 | No | Yes | N/A |
| *lgtC* | NEIS2154 | G9 | G9 | G9 | G9 | G9 | G9 | G9 | G9 | No | No | N/A |
| Uncategorised ^5^ | | | | | | | | | | | | |
| hypo. | NEIS1156 | G9 | G9 | G9 | G9 | G10 | G9 | G11 | G11 | Yes | Yes | N/A |
| hypo. | NEIS1750 | G8 | G8 | G8 | G8 | G8 | G8 | G9 | G8 | No | No | N/A |
| hypo. | NEIS2514 | C23 | C22 | C23 | C22 | C22 | C24 | C20 | C21 | Yes | Yes | N/A |
| hypo. (ATTCAG) | NEIS3139 | 12 | 12 | 12 | 12 | 12 | 12 | 12 | 12 | No | No | N/A |
| Zn-dependent protease (TTCC) | NEIS2475 | 12 | 12 | 12 | 12 | 12 | 12 | 12 | 12 | No | No | N/A |
| rubredoxin | NEIS0980 | C10 | C10 | C10 | C10 | C10 | C10 | C10 | C10 | No | No | N/A |
| putative phage replication initiation | NEIS0277 | C11 | C11 | C11 | C11 | C11 | C11 | C10 | C10 | No | Yes | Strong (6/6) |
| phosphogluconate dehydratase | NEIS1332 | C9 | C9 | C9 | C9 | C9 | C9 | C9 | C9 | No | No | N/A |
| hypo. (CAAG) | NEIS3112 | 11 | 11 | 11 | 11 | 11 | 11 | 12 | 13 | No | Yes | N/A |
| putative rotamase  (AAAGTTGCC) | NEIS0276 | 11 | 11 | 11 | 11 | 11 | 11 | 10 | 10 | No | Yes | N/A |
| (AAAGCTGCC) |  | 7 | 7 | 7 | 7 | 7 | 7 | 7 | 7 | No | No | N/A |
| lipoprotein | NEIS0009 | A11 | A11 | A11 | A11 | A11 | A11 | A11 | A11 | No | No | N/A |
| tRNA pseudouridine synthase A | NEIS2017 | C9 | C9 | C9 | C9 | C9 | C9 | C9 | C9 | No | No | N/A |
| virulence associated protein (AGCA) | NEIS2930 | 7 | 7 | 7 | 7 | 7 | 7 | 7 | 7 | No | No | N/A |
| autotransporter A, *autA* (AAGC) | NEIS1859 | 36 | 36 | 36 | 36 | 36 | 36 | 20 | 18 | No | Yes | N/A |
| iron-regulated protein *frpA*, truncation  (ATAACAAA) | NEIS3265 | 14 | 14 | 14 | 14 | 14 | 14 | 13 | 13 | No | Yes | N/A |

1: Yes – any variation in repeat number in at least one outbreak isolate.

2: Yes – consistent variation in repeat number between both nearest neighbours and 3 or more of outbreak isolates.

3: Strong – consistent variation in expression state (ON or OFF for coding PV; high, intermediate or low for IGR PV) between both nearest neighbours and ≥ five outbreak isolates. Weak – variation in expression state between both nearest neighbours and three or four outbreak isolates. Numbers in brackets indicate the number of outbreak isolates from a total of six with a consistent difference in expression states.

4: The *opa* genes are paralogous and do not have dedicated PubMLST ids for each locus. *opaABDJ* annotations assigned on the genomic context surrounding each *opa* locus as described in Dempsey *et al*., 1995.

```
igrNEIS0055_7      AAGCCAAATCCCAATATACTTATAAATGGCCTAATTATAGCACTTAATCGAAATAAATTT	60
igrNEIS0055_4      AAGCCAAATCCCAATATACTTATAAATGGCCTAATTATAGCACTTAATCGAAATAAATTT	60
                   ************************************************************

igrNEIS0055_7      ATGAGTACGTAGAGTATAATTAGTATTCTTCTTTCCAACTTCCTTATACTTATAGATTCT	120
igrNEIS0055_4      ATGAGTACGTAGAGTATAATTAGTATTCTTCTTTCCAACTTCCTTATACTTATATACTTA	120
                   ************************************************************

igrNEIS0055_7      AAAATCATG			126
igrNEIS0055_4      TAGATTCTA AAATCATG		134
                   ******
```

**Figure S2:** Clustal alignment of sequences upstream of the *cssA* open reading frame: ATG, start codon in red; yellow and green highlights one or two copies of TATACTTA, respectively; bold, predicted ribosome binding site. Allele 7 is present in all outbreak variants and closest relatives.

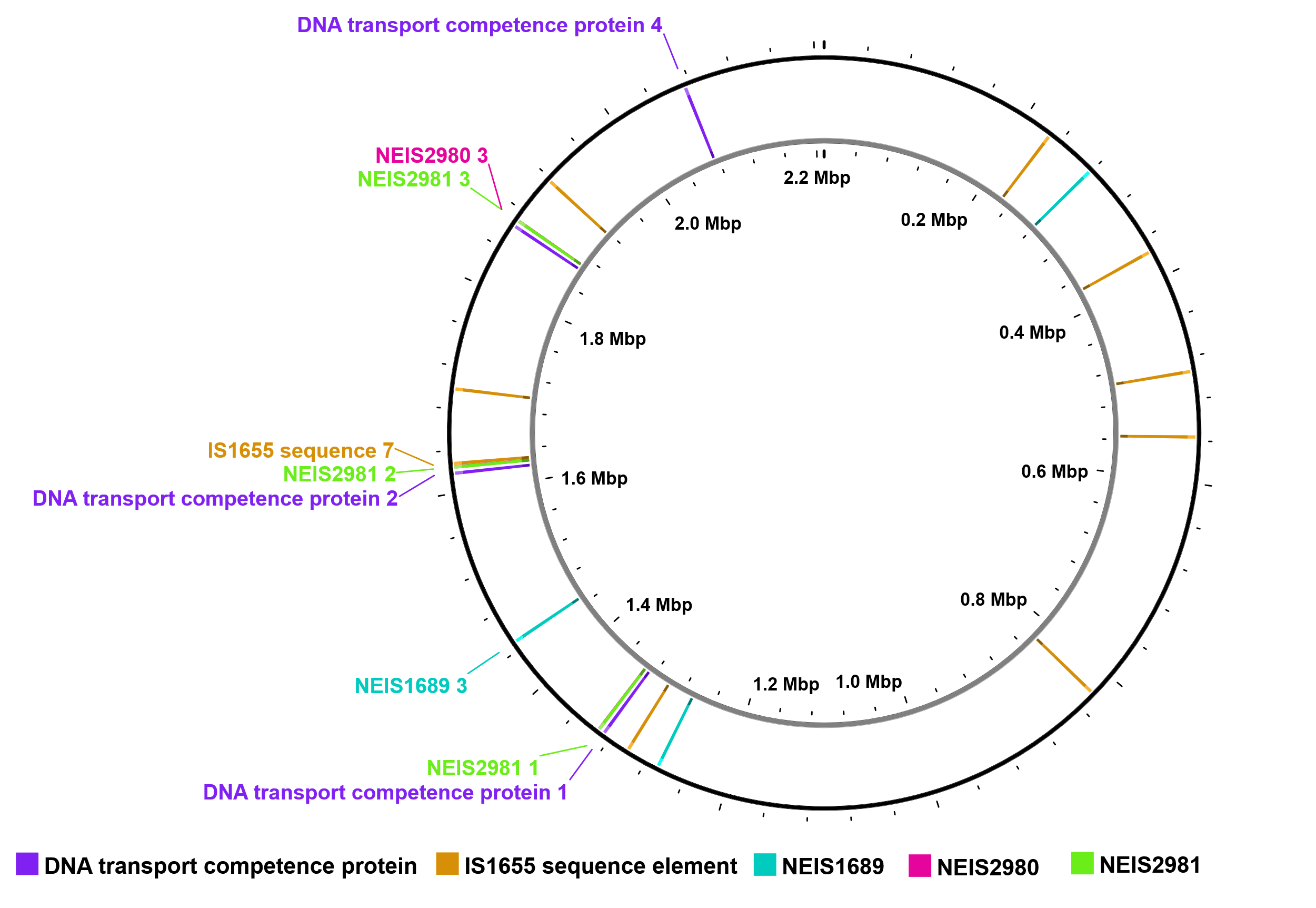

**Figure S3**: Paralogous loci mapped to the 193899 genome. Labels indicate loci with variation between the Kent outbreak variant (PubMLST id:193899) and their closest relatives (PubMLST ids: 194251 and 194252). DNA transport competence protein 4 is also variable among the six outbreak isolates.

NEIS 1464 SecA

```
Alignments:

99.8% identity in 916 residues overlap; Score: 4683.0; Gap frequency: 0.0%

Sequence1 is allele 3732; sequence 2 is allele 34

Sequence1      1 MLTNIAKKIFGSRNDRLLKQYRKSVARINALEEQMQALSDADLQAKTAEFKQRLADGQTL
Sequence2      1 MLTNIAKKIFGSRNDRLLKQYRKSVARINALEEQMQALSDADLQAKTAEFKQRLADGQTL
                 ************************************************************

Sequence1     61 DGILPEAFAVCREASRRTLGMRHFDVQLIGGMVLHDGKIAEMRTGEGKTLVATLAVYLNA
Sequence2     61 DGILPEAFAVCREASRRTLGMRHFDVQLIGGMVLHDGKIAEMRTGEGKTLVATLAVYLNA
                 ************************************************************

Sequence1    121 LAGKGVHVVTVNDYLASRDAGIMEPLYNFLGLTVGVIISDMQPFDRQNAYAADITYGTNN
Sequence2    121 LAGKGVHVVTVNDYLASRDAGIMEPLYNFLGLTVGVIISDMQPFDRQNAYAADITYGTNN
                 ************************************************************

Sequence1    181 EFGFDYLRDNMVTDQYDKVQRELNFAVVDEVDSILIDEARTPLIISGQADDNIQLYQIMN
Sequence2    181 EFGFDYLRDNMVTDQYDKVQRELNFAVVDEVDSILIDEARTPLIISGQADDNIQLYQIMN
                 ************************************************************

Sequence1    241 TVPPHLVRQETEEGEGDYWVDEKAHQVILSEAGHEHAEQILTQMGLLAENDSLYSAANIA
Sequence2    241 TVPPHLVRQETEEGEGDYWVDEKAHQVILSEAGHEHAEQILTQMGLLAENDSLYSAANIA
                 ************************************************************

Sequence1    301 LMHHLMAALRAHSLFHKDQHYVIQDGEIVIVDEFTGRLMSGRRWSEGLHQAVEAKEGVEI
Sequence2    301 LMHHLMAALRAHSLFHKDQHYVIQDGEIVIVDEFTGRLMSGRRWSEGLHQAVEAKEGVEI
                 ************************************************************

Sequence1    361 KRENQTLASITFQNYFRLYTKLSGMTGTADTEAFEFQSIYNLETVIIPTNRPVQRKDFND
Sequence2    361 KRENQTLASITFQNYFRLYTKLSGMTGTADTEAFEFQSIYNLETVIIPTNRPVQRKDFND
                 ************************************************************

Sequence1    421 QIFRSAEEKFEAVVKDIEECHKRGQPVLVGTTSIENSELVSHLLQKAGLPHNVLNAKEHE
Sequence2    421 QIFRSAEEKFEAVVKDIEECHKRGQPVLVGTTSIENSELVSHLLQKAGLPHNVLNAKEHE
                 ************************************************************

Sequence1    481 REALIVAQAGKVGAITVATNMAGRGTDIVLGGNLKHQTDAIRADEALSDEEKQAQIAALE
Sequence2    481 REALIVAQAGKVGAITVATNMAGRGTDIVLGGNLKHQTDAIRADETLSDEEKQAQIAALE
                 ********************************************* **************

Sequence1    541 NGWQAEHDKVMEAGGLHIIGTERHESRRIDNQLRGRSGRQGDPGSSRFYLSFEDPLLRLF
Sequence2    541 NGWQAEHDKVMEAGGLHIIGTERHESRRIDNQLRGRSGRQGDPGSSRFYLSFEDPLLRLF
                 ************************************************************

Sequence1    601 ALDRAAAILNRLAPERGVAIEHNLLTRQIEGAQRKVEGRNFDMRKQVLEYDDVANEQRKV
Sequence2    601 ALDRAAAILNRLAPERGVAIEHNLLTRQIEGAQRKVEGRNFDMRKQVLEYDDVANEQRKV
                 ************************************************************

Sequence1    661 IYSQRNEILTSKDISDLMKEIRSDVVSDLVDTYMPPDSMEEQWDIPTLENRLAAEFRLHE
Sequence2    661 IYSQRNEILTSKDISDLMQEIRSDVVSDLVDTYMPPDSMEEQWDIPTLENRLAAEFRLHE
                 ****************** *****************************************

Sequence1    721 DIQSWLKADNAIDGQDIKERLIERIENEYAAKTELVGKQAMADFERNVMLQVIDNQWREH
Sequence2    721 DIQSWLKADNAIDGQDIKERLIERIENEYAAKTELVGKQAMADFERNVMLQVIDNQWREH
                 ************************************************************

Sequence1    781 LAAMDYLRQGIHLRSYAQKNPKQEYKREAFTMFQDLWNGIKFHIASLLTSVQIEQNPVAV
Sequence2    781 LAAMDYLRQGIHLRSYAQKNPKQEYKREAFTMFQDLWNGIKFHIASLLTSVQIEQNPVAV
                 ************************************************************

Sequence1    841 VEEQPIGNIQSIHSESPDMEELLGQSQTDLVTEAFNPDGTDFSPEALEARGQIVHRNDPC
Sequence2    841 VEEQPIGNIQSIHSESPDMEELLGQSQTDLVTEAFNPDGTDFSPEALEARGQIVHRNDPC
                 ************************************************************

Sequence1    901 PCGSGLKYKQCHGKLA
Sequence2    901 PCGSGLKYKQCHGKLA
                 ****************
```

**Figure S4:** Alignment of the alleles for NEIS1464 (preprotein translocase subunit *secA*). There are eight SNVs between allele 3732 and closest relative’s allele 34, located in the latter part of the sequence from nt. 1570 onwards, with NS SNVs highlighted in yellow.

**Table S5** Pathway enrichment analysis comparing the closest relative (PubMLST id 194252) to the outbreak variant (PubMLST id 193899). All significant ontology terms (q<0.05) under any of the three annotation routes. “scheme” is the term vocabulary (GO, KEGG, COG); “source” is how the annotation was obtained – transfer from the MC58 reference, profile search against the COG database, or orthology assignment by eggNOG-mapper. “k” is the number of genes carrying the term among those that differed, “K” the number carrying it in the 2008-gene background, and “fold” the resulting enrichment.

| **term** | **scheme** | **source** | **description** | **k/K** | **fold** | **q** |
| --- | --- | --- | --- | --- | --- | --- |
| N | COG | profile | Cell motility | 6/20 | 13.1 | 2e-05 |
| N | COG | orthology | Cell motility | 6/20 | 12.4 | 3.8e-05 |
| U | COG | orthology | Intracellular trafficking and secretion | 8/60 | 5.5 | 0.00027 |
| GO:0015627 | GO | transfer | type II protein secretion system complex | 3/3 | 39.1 | 0.00028 |
| GO:0015628 | GO | transfer | protein secretion by the type II secretion | 3/4 | 29.3 | 0.00055 |
| GO:0016651 | GO | transfer | oxidoreductase activity, acting on NAD(P)H | 2/2 | 39.1 | 0.0038 |

**Table S7** Loci where the outbreak (OB) and closest available relative (CR)have different alleles in both the Kent and Southampton outbreaks. Bold indicates genes predicted to be different between OB and NN strains through HGT events.  New, new alleles; ?, partial sequence

| **Gene** | **Kent** | | **Southampton** | |
| --- | --- | --- | --- | --- |
|  | **OB** | **CR** | **OB** | **CR** |
| NEIS0041  competence protein | 71 | 325 | 33 | awaited |
| NEIS0202 | **44** | **5** | 2 or 86 | 36 |
| NEIS0210  PilE | 2223 | ? | 4 | 32 |
| NEIS0213  glycosyl transferase | new | 288 | 54 | ? |
| NEIS0380  acetyl transferase | unique |  | **28** | **684** |
| NEIS0400  glycosyl transferase | unique | 101 | 55 | 37 |
| NEIS0826 Replicative helicase | **5773** | **101** | **148** | **2** |
| NEIS0827  PilH | **55** | **41** | **new** | **35** |
| NEIS0828  PilI | **13** | **36** | **48** | **1** |
| NEIS0829  PilJ | **310** | **42** | **112** | **3** |
| NEIS0839  PilK | **245** | **31** | **103** | **2** |
| NEIS0831  PilX | **2053** | **33** | **103** | **2** |
| NEIS0955 hypo | **146** | **new** | **1** | **?** |
| NEIS0956 cell surface protein | **new** | **new** | **7** | **?** |
| NEIS1156 | 137 | new | 1 | ? |
| NEIS1194 NgoAX methylase | new | 152, 40 | various | ? |
| NEIS1310 NgoAXII methylase | 987 | 381 | 6 or 7 | 16 |
| NEIS1403  Opa | 1031 | 135 | 231/new | ? |
| NEIS1880  DNA transport protein | 740 | 308 | 22 | ? |
| NEIS1996  DNA competence protein | 747 | 308 | 23 | ? |
| NEIS2020  PorB | 10381 | 42 | 192 | 1 |
| NEIS2575  C term  Toxicity domain | new | new | new | new |
| NEIS2649 | new | new | new | new |
| NEIS3112 | new | new | new | new |
| NEIS3229 | new | new | new | new |
| NEISp0276   SurA PPIase | 386 | 13 | 31; 60 | 31 |
